# Evolutionary dynamics of the OR gene repertoire in teleost fishes: evidence of an association with changes in olfactory epithelium shape

**DOI:** 10.1101/2021.03.09.434524

**Authors:** Maxime Policarpo, Katherine E Bemis, James C Tyler, Cushla J Metcalfe, Patrick Laurenti, Jean-Christophe Sandoz, Sylvie Rétaux, Didier Casane

## Abstract

Teleost fishes perceive their environment through a range of sensory modalities, among which olfaction often plays an important role. Richness of the olfactory repertoire depends on the diversity of receptors coded by homologous genes classified into four families: OR, TAAR, VR1 and VR2. Herein, we focus on the OR gene repertoire. While independent large contractions of the OR gene repertoire associated with ecological transitions have been found in mammals, little is known about the diversity of the OR gene repertoire and its evolution within teleost fishes, a group that includes more than 34,000 living species. We analyzed genomes of 163 species representing diversity in this large group. We found a large range of variation in the number of functional OR genes, from 15 in *Syngnathus typhle* and *Mola mola*, to 429 in *Mastacembelus armatus*. The number of OR genes was higher in species with an extensively folded olfactory epithelium, that is, for example, when a multi-lamellar rosette was present in the olfactory organ. Moreover, the number of lamellae was correlated with the richness of the OR gene repertoire. While a slow and balanced birth-and-death process generally drives evolution of the OR gene repertoire, we inferred several episodes of high rates of gene loss, sometimes followed by large gains in the number of OR genes. These gains coincide with morphological changes of the olfactory organ and suggest a strong functional association between changes in the morphology and the evolution of the OR gene repertoire.

## Introduction

Olfaction is an important sensory modality in many animals because it serves essential functions such as feeding, reproduction, migration, kin recognition, and predator avoidance. In vertebrates, odorant molecules are primarily detected by olfactory sensory neurons, each expressing an olfactory receptor, following the one neuron - one receptor rule, although some exceptions have been reported (Sato, et al. 2007). Olfactory receptors are G protein-coupled receptors coded by a large gene family (Buck and Axel 1991). The diversity of odors that an individual can discriminate depends on the richness of its olfactory gene repertoire, which results from the number and diversity of functional genes present in its genome. There is a great range of variation in the number and diversity of olfactory receptor genes among species (Nei, et al. 2008; Niimura 2009; Khan, et al. 2015; Sharma, et al. 2019), which may reflect differences in the importance of olfaction relative to other sensory systems. Expansion and contraction of the olfactory receptor gene repertoire in a lineage is governed by a birth-and-death process; that is, recurrent gene duplications and losses that are involved in both the adaptive and non-adaptive evolution of olfaction (Nei, et al. 2008).

Vertebrate olfactory receptors genes are coded by four large multigene families named OR (olfactory receptor), TAAR (trace amine-associated receptor), V1R and V2R (vomeronasal receptor type 1 and 2) (Nei, et al. 2008). OR and TAAR genes share sequence similarity and have no introns in the coding sequence. Studies of the evolution of the vertebrate olfactory gene repertoire have focused mainly on mammalian OR genes, and far less research has been undertaken on the OR genes of bony fishes (Osteichthyes). The average number of functional OR genes is about 800 in mammals, but the range of variation is very large, from 58 in the Common Bottlenose Dolphin *Tursiops truncatus* to 2,514 in the African Elephant *Loxodonta africana* (Hughes, et al. 2018). In teleosts, the average number of OR genes is an order of magnitude smaller. Although few species of teleosts have been analyzed in prior studies, a wide range of variation has been observed (Niimura 2009; Jiang, et al. 2019).

Vertebrate OR genes are classified into two types and eleven families that diverged before the separation of actinopterygians and sarcopterygians: Type 1 (α, β, γ, δ, ε and ζ) and Type 2 (η, θ1, θ2, κ and λ). There is a clear separation between Type 1 and Type 2 genes, but the phylogenetic relationships among the gene families belonging to each type are not well established (Niimura and Nei 2005; Niimura 2009). In tetrapods, the γ family is greatly expanded and, to a lesser extent, the α family also is expanded. Sizes of the other gene families decreased and some disappeared in some lineages. In amniotes, most OR genes belong to the α or γ families. In contrast, teleosts lack α genes and only one γ gene exists in some species, but all other gene families are present (Lv, et al. 2019). These observations led to the hypothesis that α and γ genes serve to detect airborne odorants while other gene families detect water-soluble odorants (Niimura and Nei 2005). Hence, gene families would have expanded or contracted differentially after the transition from an ancestral aquatic and aerial olfactory system, in the last common ancestor of bony fishes, to either only aerial olfaction in tetrapods or only aquatic olfaction in teleosts. At smaller phylogenetic scales, high rates of OR gene death have been linked to major ecological transitions. For example, cetaceans lost many OR genes because of their return to the sea (Liu, et al. 2019). In primates, an acceleration of OR gene loss that occurred in the ancestral branch of haplorhines is associated with the acquisition of acute vision, and a high rate of OR gene loss was also observed at the ancestral branch of leaf-eating colobines related with the dietary transition from frugivory to folivory (Niimura, et al. 2018). Conversely, an extreme expansion of the OR gene repertoire occurred in elephants, although the reason for this is unknown (Niimura, et al. 2014). A recent study analyzed how the OR gene repertoire has undergone expansion and contraction across Mammalia with respect to several ecological adaptations (Hughes, et al. 2018). Another study found a correlation between the number of OR genes and the relative size of the cribriform plate, a part of the ethmoid bone, and the area of its foramina, which are indicators of the relative number of olfactory sensory neurons (Bird, et al. 2018).

In contrast, we know little about the evolution of OR in teleosts. With > 34,000 species, teleosts represents about half of the extant vertebrate species and more than 95% of aquatic vertebrates and are an ideal model to study evolution of the olfactory system in aquatic environments. Molecular datings suggest an origin of teleosts as far back as the Late Carboniferous/Early Permian (Near, et al. 2012), although fossils clearly assignable to Teleostei do not appear until the Mesozoic. This vast period of time allowed great diversification of morphology, physiology, behavior, ecology and habitat. Switches in sensory modalities have involved large numbers of gene births and deaths in teleosts. For example, several deep-sea fishes independently evolved rod opsin-based dim light vision relying on the expansion of RH1 gene repertoires (Musilova, et al. 2019). In some cavefishes living in total darkness, loss of vision allowed the decay of many eye-specific genes (Policarpo, et al. 2020). Much less is known about the evolution of olfaction and the dynamics of the OR gene repertoire size.

Since the seminal work of R. H. Burne at the beginning of the 20th century (Burne 1909), the relative importance of olfaction among sensory systems in teleosts has been assessed by studying the shape and organization of the olfactory epithelium (OE). Most teleosts have multi-lamellar OE (ML-OE) that forms a rosette, with each lamella increasing the epithelial surface, suggesting that the olfactory system is well-developed (Kasumyan 2004). The presence of a rosette may be the ancestral state for teleosts (Hansen and Zielinski 2005), indicating that the sense of smell was well developed in their last common ancestor. However, simpler organizations have been observed in many species. For example, the OE is flat (no lamellae) in Broad-nose Pipefish *Syngnathus typhle (Dymek, et al. 2020)* and there is only one lamella in the Round Goby *Neogobius melanostomus (Hansen and Zielinski 2005)*. When a rosette is present, the number of lamellae is highly variable, with up to 230 in the Barred Pargo *Hoplopagrus guentherii* (Hara 1975). Species with a simple OE, such as the Three-spined Stickleback *Gasterosteus aculeatus*, the Crowned Seahorse *Hippocampus coronatus*, the Green Pufferfish *Dichotomyctere fluviatilis* are assumed to have degenerate olfaction. Conversely, species such as the Japanese Eel *Anguilla japonica*, the Zig-zag Eel *Mastacembelus armatus*, the Channel Catfish *Ictalurus punctatus* and the Catla *Labeo catla* that possess rosettes with many lamellae may rely primarily on olfaction to perceive important features of their environments. Although many teleost genomes are now available, correlation between the complexity of the olfactory organ and richness of the OR gene repertoire has not been investigated. To date, OR gene repertoires of teleosts have been reported for only a handful of species (Niimura and Nei 2005; Niimura 2009; Gao, et al. 2017; Jiang, et al. 2019; Lv, et al. 2019), and no comparative analyses have been made.

The purpose of our study was to describe the evolutionary dynamics of the OR gene repertoire across the diversity of teleosts. We conducted large-scale phylogenetic analyses of the OR gene repertoires of 163 species to examine correlations between contractions and expansions of the OR gene family and changes in morphological traits, ecological parameters, and genome size. We found a positive correlation between the richness of the OR gene repertoire and both the presence of a multi-lamellar olfactory epithelium and the number of lamellae. In particular, we discovered that the two highest death rates of OR genes coincide with two independent transitions from a multi-lamellar OE to a simple OE. Moreover, a re-acquisition of a multi-lamellar OE coincides with a re-expansion of the OR repertoire. These observations suggest a strong functional link between changes in the morphology of the olfactory organ and the evolution of the OR gene repertoire.

## Results

### Diversity and evolution of OR gene repertoire in teleost fishes

The genomes of 307 teleost species and two non-teleostean actinopterygians (Sterlet Sturgeon *Acipenser ruthenus* and Spotted Gar *Lepisosteus oculatus*) were downloaded from NCBI and ENSEMBL databases.

We discarded six genomes for which the genome assembly size was not similar to the genome size estimated with other methods and compiled in the Animal Genome Size Database (Gregory 2020) (**fig. 1A** and **supplementary fig. S1, Supplementary Material** online). The completeness of the remaining 301 genomes was assessed using BUSCO (Waterhouse, et al. 2018). We discarded all genomes for which < 90% of the BUSCO genes were retrieved as complete coding sequences. This threshold allowed us to select a set of 163 teleost and two non-teleost genomes for which we could expect to identify a nearly complete set of OR genes. A higher threshold would have removed more species from consideration and would not have greatly improved the average quality of the genomes analyzed (**fig. 1B**).

**Fig. 1.**
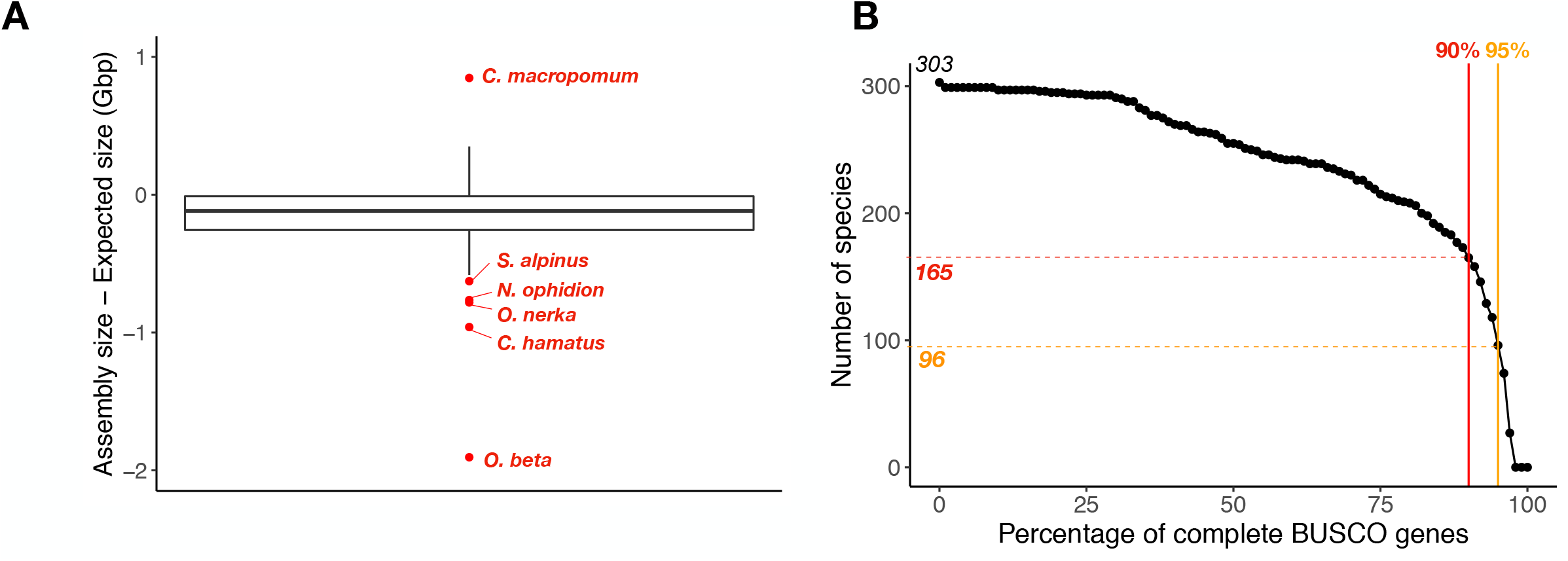
Genome selection. (A) Congruence of genome assembly size and expected genome size based on haploid DNA content (C−value x 0.978 × 10^9^) for 309 actinopterygian fishes (307 teleosts and 2 non-teleosts); six incongruent estimates of genome size are shown in red and species names indicated (more details in **supplementary fig. S3, Supplementary Material**). (B) Number of genomes according to a completeness threshold (% of complete BUSCO genes). Out of 303 genomes with congruent estimates of genome size, 165 (163 teleosts and 2 non-teleosts) show a completeness > 90%, and 96 genomes (96 teleosts) a completeness > 95%.

We found 27,346 OR genes in the genomes of these 165 species. Among the OR genes, 19,270 were complete with seven transmembrane domains and we refer to these as functional olfactory receptors; 4,869 genes carried at least one frameshift or a stop codon (also called loss-of-function (LoF) mutations) and were classified as pseudogenes. We also identified 2,347 truncated genes, which were incomplete sequences without any loss-of-function mutations, and 860 edge genes which were incomplete coding sequences located at scaffold edges (**fig. 2A**).

**Fig. 2.**
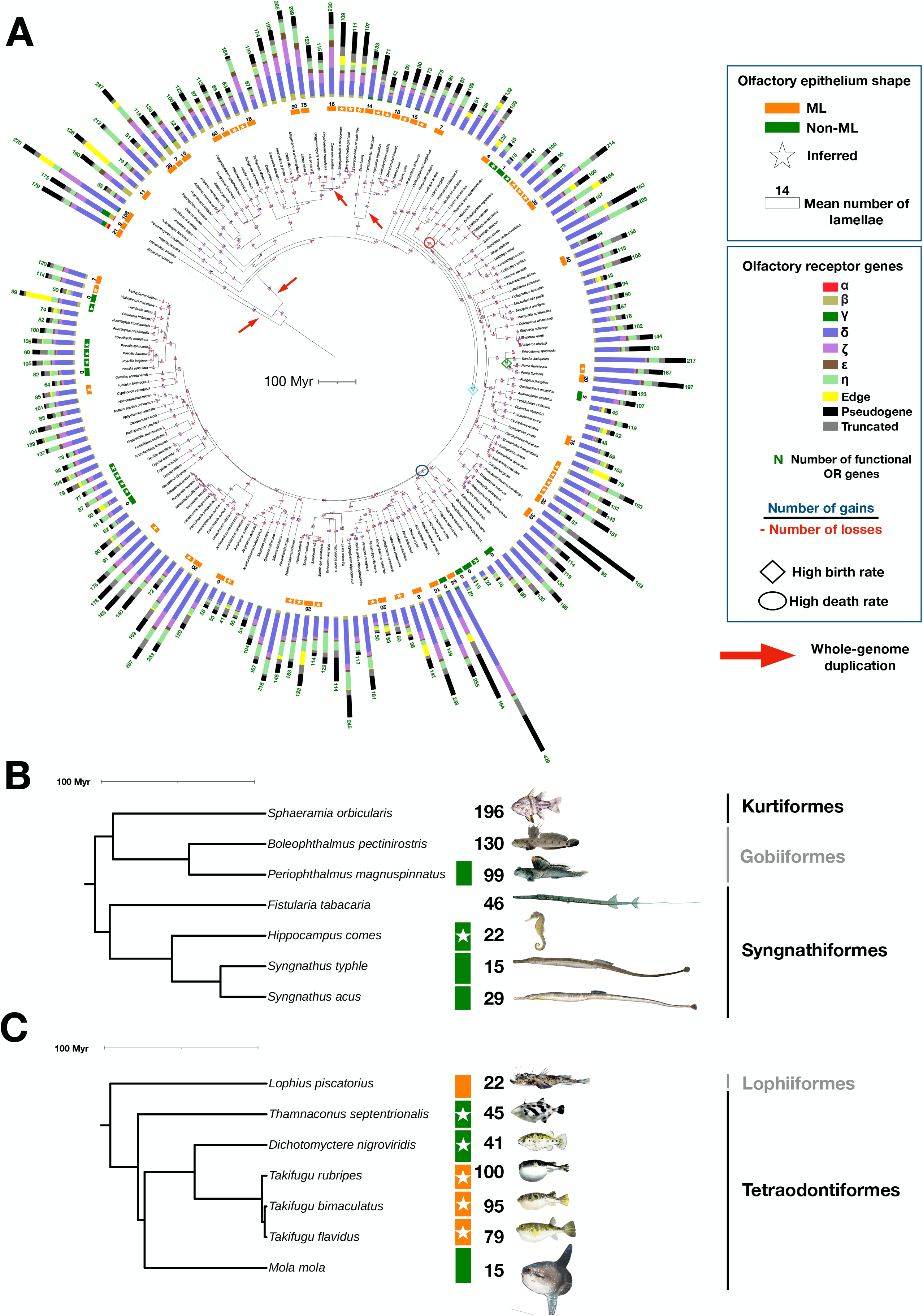
Chronograms. (A) Dated phylogeny for 163 teleost fishes and two outgroups (from https://fishtreeoflife.org/). For each species, the total number of functional OR genes, the proportion of genes from each gene family, the proportion of truncated and edge genes, and the proportion of pseudogenes are provided. When available from the literature, presence of a multi-lamellar olfactory epithelium (ML-OE) or a simple olfactory epithelium (Non-ML-OE) is reported. If found in the literature, the number of lamellae in ML-OE is also indicated. Presence or absence of a ML-OE was inferred for some species without data on the shape of the olfactory epithelium using information available for a congeneric species (white star); the number of lamellae was not inferred because of intrageneric variation. Inferred number of gene gains and losses are provided on each branch (corresponding birth and death rates are given in **supplementary fig. 7, Supplementary Material**). The branches associated with the two highest birth rates and the two highest death rates are indicated by diamond and oval symbols, respectively. Whole-genome duplications are indicated by red arrows. The tree was annotated and visualized using iTOL. (B) Focus on Syngnathiformes and related species. (C) Focus on Tetraodontiformes and related species.

The number of truncated genes per genome was correlated with the number of pseudogenes (r = 0.62, p-value = 2.2e-16), whereas the number of edge genes was not correlated with the number of pseudogenes (r = 0.045, p-value = 0.57), or with the number of truncated genes (r = 0.031, p-value = 0.7) (respectively **supplementary fig. S2A, B and C, Supplementary Material** online). These results suggest that a large fraction of truncated genes are due to large deletions and correspond to actual pseudogenes, whereas edge genes are mainly assembly artifacts.

We found a positive correlation between the number of pseudogenes and the number of functional genes (r = 0.42, p-value = 3e-08) (**supplementary fig. S3A, Supplementary Material** online). The phylogenetic signal (*i*.*e*., the tendency of closely related species to resemble each other more than species drawn at random from the same phylogenetic tree) for both the number of functional ORs and the number of pseudogenes was strong (Pagel’s λ = 0.77 and 0.93, respectively, and both significantly different from 0). Therefore, we applied a phylogenetic correction, after which the correlation still holds true (r = 0.36, p-value = 2.5e-6) (**supplementary fig. S3B, Supplementary Material** online). This suggests that OR repertoire remodeling is a slow process. Conversely, there was no correlation between the proportion of pseudogenes and the number of functional genes (r = 0.013, p = 0.86) (**supplementary fig. S2D, Supplementary Material** online), suggesting that the rate of gene loss does not increase with the number of functional genes. These results are similar to those obtained with the OR genes of placental mammals (Niimura, et al. 2014).

We assessed the relative efficacy of our pipeline to identify the OR gene repertoire in a genome, including both functional and non-functional sequences, by comparing the number of genes retrieved in the present study and in previous analyses of the OR repertoire in teleosts, that is *Danio rerio, Gasterosteus aculeatus, Oryzias latipes, Takifugu rubripes Dichotomyctere nigroviridis* (Niimura 2009), *Ictalurus punctatus* (Gao, et al. 2017), *Siniperca chuatsi* (Lv, et al. 2019), and *Pseudoliparis swirei* (Jiang, et al. 2019). We systematically found more OR genes than in these previous reports, and, for several species, many more functional sequences (**table 1**). This is probably because of the higher quality of more recent genome versions when compared with older versions used in previous studies. This may also result from a better pipeline implementation compared to other analyses based on the same genome versions (more details are given in **supplementary fig. S4A-H, Supplementary Material** online).

**Table 1.**
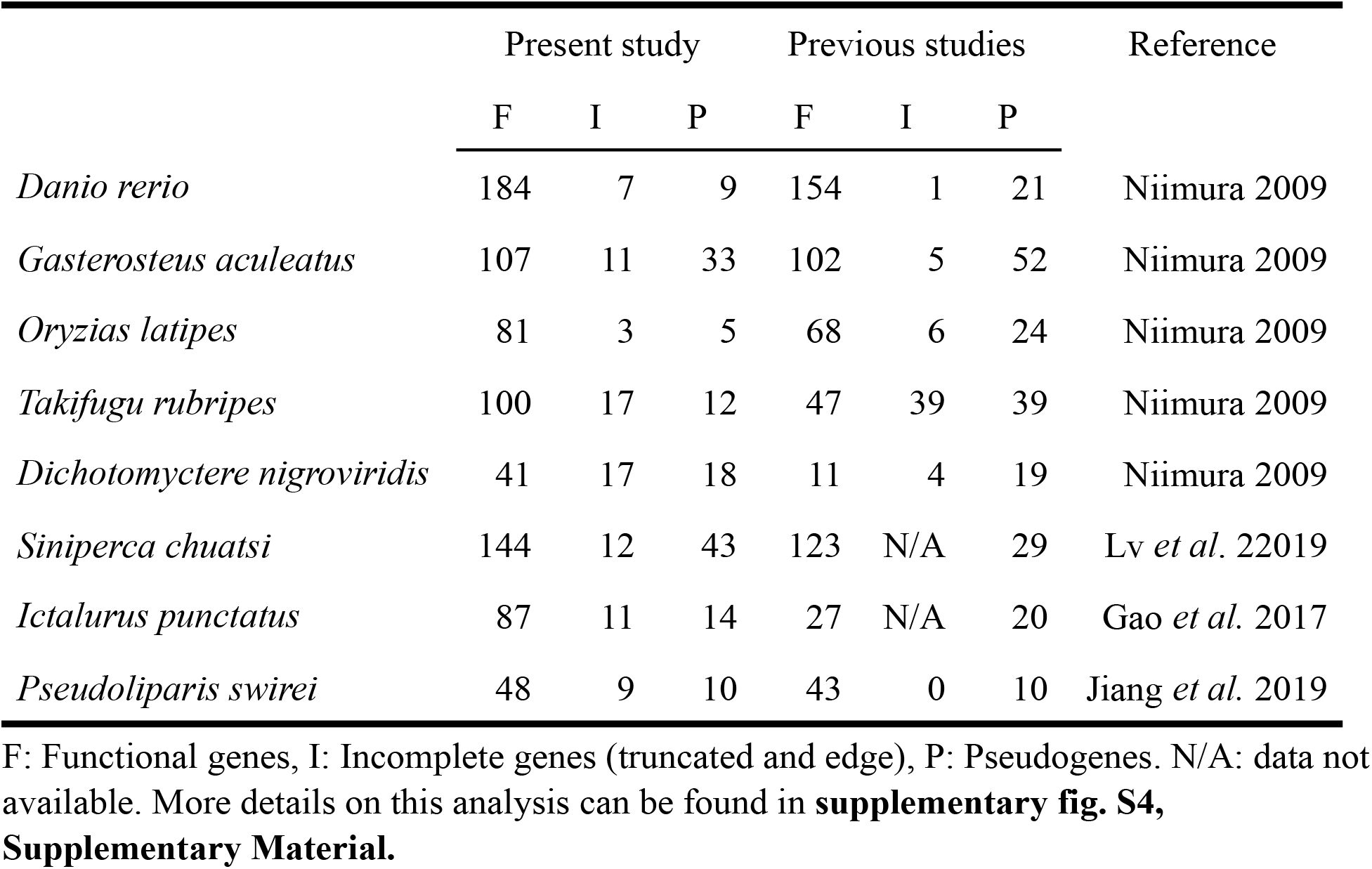
Comparison of the number of OR genes retrieved from the genome of eight species of teleosts in the present and previous studies.

OR genes were classified into seven previously defined families (Niimura and Nei 2005): α, β, γ, ζ, ε, δ (Type 1) and η (Type 2). Genes of the α family are absent in teleosts, but they are present in the two non-teleost outgroups, *Acipenser ruthenus* and *Lepisosteus oculatus* (**fig. 2A**). Type 2 genes belonging to the families θ, κ and λ were not analyzed because they may not be OR genes, most often present as a unique copy in vertebrates when they are not absent, with very rare cases of gene gains and losses (Niimura 2009).

To visualize gene sequence diversity within each gene family, we reconstructed a phylogenetic tree with the functional OR sequences of 38 species belonging to 38 teleost orders, each species having the most complete genome within that order. At the scale of teleosts, this phylogeny (**supplementary fig. S5, Supplementary Material** online) shows that the δ family contains many more genes and is much more diverse than the other families, followed by the ζ and η families. The β and ε families are much less diverse, and diversity is very low in the γ family. This observation also holds at the species level (**fig. 2A**).

### Contrasting richness of the OR repertoire among teleost fishes

We found an unanticipated large range of variation in the number of OR genes among teleosts (**fig. 2, supplementary fig. S6, Supplementary Material** online). The mean number of functional OR genes per species was 117, with the smallest number (15) in *Syngnathus typhle* (Syngnathiformes) and *Mola mola* (Tetraodontiformes) and the highest (429) in *Mastacembelus armatus* (Synbranchiformes). The other Syngnathiformes species studied (*Syngnathus acus* and *Hippocampus comes*) were also OR-poor, and only a few δ and η genes were found (**fig. 2B**). In Tetraodontiformes, including *Mola mola, Thamnaconus septentrionalis, Dichotomyctere nigroviridis* and three *Takifugu* species, the number of OR genes was more variable (**fig. 2C**).

The number of gene gains and losses along each branch of the phylogenetic tree was inferred using the gene tree - species tree reconciliation method (Chen, et al. 2000), with one gene tree per OR gene family (**fig. 2A**). Birth and death rates were computed using these inferred numbers of gene gains and losses (**supplementary fig. S7, Supplementary Material** online). The mean birth and death rates were very similar, 0.0071 and 0.0073 per gene per million years, respectively (**fig. 3**), but the death rate was particularly high in the branches leading to Tetraodontiformes (0.3) and Syngnathiformes (0.32).

**Fig. 3.**
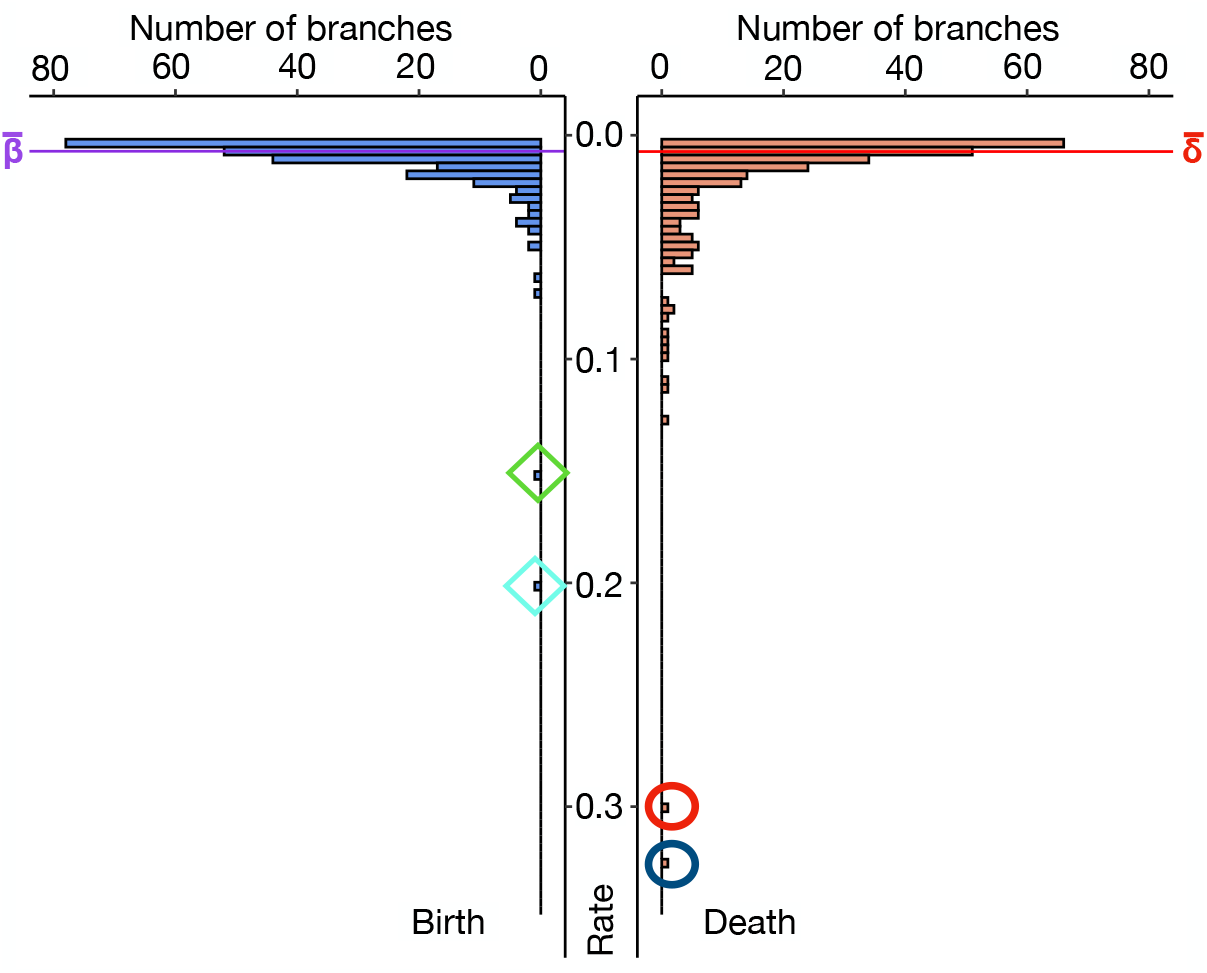
Distribution of birth and death rates along the branches of the phylogenetic tree shown in **fig. 2A**. The mean values are indicated and the two highest birth rates and the two highest death rates are highlighted with diamond and oval frames, respectively.

We also found two cases of high birth rates. The highest birth rate (0.2) was at the level of a deep internal node, in the common ancestor of Labriformes and Cyprinodontiformes while the second highest birth rate (0.15) was observed in the common ancestor of *Perca* + *Sander* (**fig. 2, supplementary fig. S7, Supplementary Material** online).

The number of gene losses inferred in terminal branches correlated with the sum of the number of pseudogenes and truncated genes (**fig. 4**). Interestingly, the correlation was higher when considering only the shortest terminal branches (R > 0.7 with maximum branch length < 7 Myr) and decreased progressively when longer terminal branches were added (R = 0.17 with all terminal branches, see also **supplementary fig. S8, Supplementary Material** online). Similar results were obtained using either the number of pseudogenes or the number of truncated genes, but no correlation was found with edge genes (**supplementary fig. S9, Supplementary Material** online).

**Fig. 4.**
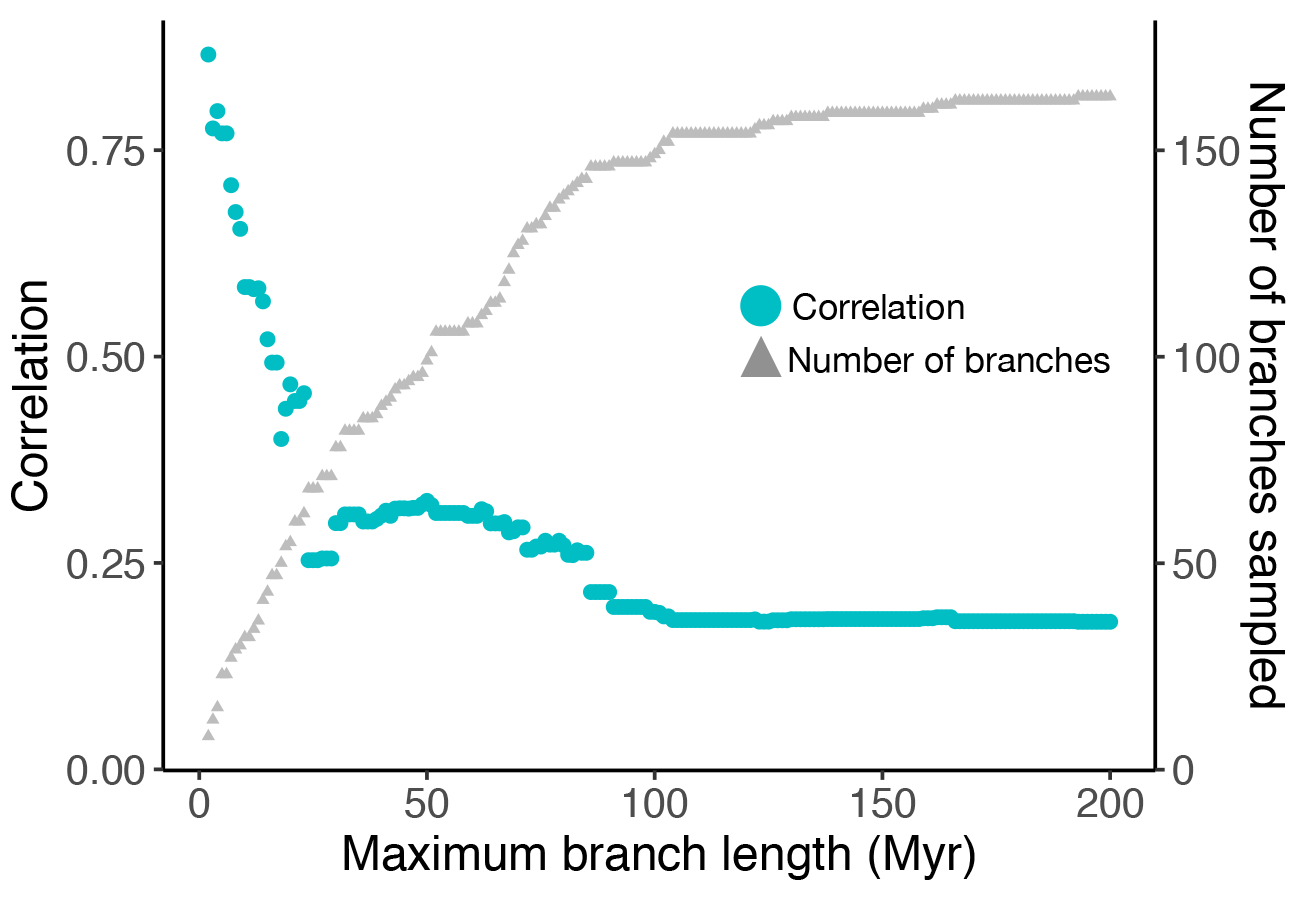
Variation in the correlation between the number of gene losses inferred using a gene tree - species tree reconciliation method and the number of observed pseudogenes and truncated genes in the external branches of the phylogenetic tree according to a maximum branch length threshold ranging from 2 to 200 Myr. For each threshold, the number of branches is indicated by a gray triangle.

We ran simulations of sequence evolution involving nucleotide substitutions and small indels taking into account their relative frequencies estimated in a study of gene decay in teleosts (Policarpo, et al. 2020). We found that about half of the pseudogenes were not detected after ∼ 40 millions of generations (**supplementary fig. S10, Supplementary Material** online). If we also consider that large deletions can remove large fragments of non-functional genes and even complete sequences, then the time required for a large fraction of pseudogenes to fade away is probably much shorter. The continuous process of divergence and removal of non-functional genes may explain why the expected correlation between the inferred number of gene losses and the observed number of pseudogenes and truncated genes is high only when short terminal branches are analyzed.

### Evolution of olfactory epithelium folding in teleost fishes

Using the most recent hypotheses about teleost phylogeny and a compilation of descriptions on the organization of the olfactory organ, we inferred the ancestral state of the OE and its folding using maximum likelihood and parsimony methods. We conducted a literature survey of the shape of the OE for 220 actinopterygians beginning with the work of R. H. Burne (Burne 1909) (**supplementary Data S1, Supplementary Material** online). Our analysis indicated that a multi-lamellae rosette is ancestral, which was expected (Hansen and Zielinski 2005), but we also refined aspects of its evolution. In particular, the ML-OE has been lost several times and regained in some lineages (**fig. 2, supplementary fig. S11, Supplementary Material** online). Maximum likelihood and parsimony methods gave very similar results, except for a few internal nodes, such as the most recent common ancestor of Syngnathiformes and Gobiiformes which is inferred to have a ML-OE with the maximum likelihood MPPA and parsimony with DELTRAN (**supplementary fig. S11 A and C, Supplementary Material** online), undefined with DOWNPASS (**supplementary fig. S11 B, Supplementary Material** online) but inferred to have a non-ML OE with ACCTRAN (**supplementary fig. S11 D, Supplementary Material** online).

### Correlation of the size of the OR repertoire with morphological and ecological traits

We examined the possible correlation between folding of the olfactory epithelium (evaluated as the presence or absence of a ML-OE), the extent of folding of the OE when present (evaluated as the number of lamellae), and the size of the OR gene repertoire.

Among the 220 species for which we had information about OE folding, we had estimates of the size of the OR gene repertoire for 41 species. For 37 of the 41 species we found information about the mean number of lamellae per OE. For 38 additional species for which we had an estimate of the size of the OR gene repertoire, we inferred OE shape using information available for at least one species in the same genus (**fig. 2, supplementary Data S1, Supplementary Material** online), assuming that closely related species have an OE with a similar shape (presence or absence of lamellae). However, we did not use information about the number of lamellae in a given species to infer this character at the genus scale because it is much more labile (Kasumyan 2004).

Olfactory epithelium shape was coded as a binary character: (1) for a multi-lamellar OE (ML-OE) with more than 2 lamellae; (0) for a flat OE or an OE with only 1 or 2 lamellae, the surface usually without ornamentation, but sometimes indented with pits (collectively called non-ML-OE).

Species with a non-ML-OE had significantly fewer functional OR genes than species with a ML-OE (**fig. 5A-B**). Olfactory epithelium shape had a strong phylogenetic signal (Pagel’s λ = 0.999, significantly different from 0). To account for the phylogenetic signal, we used a phylogenetic generalized linear model; the correlation between the OE shape and the number of OR genes was still significant (**fig. 5A-B** - **supplementary fig. S12, Supplementary Material** online). This correlation was statistically significant with the whole data set (p-value = 0.034, 79 species, including the 38 species for which we inferred OE shape, **fig. 5A**) and significant with the reduced data set containing only 41 species for which OE shape was directly known from the literature (p-value = 0.025, **fig. 5B**).

**Fig. 5.**
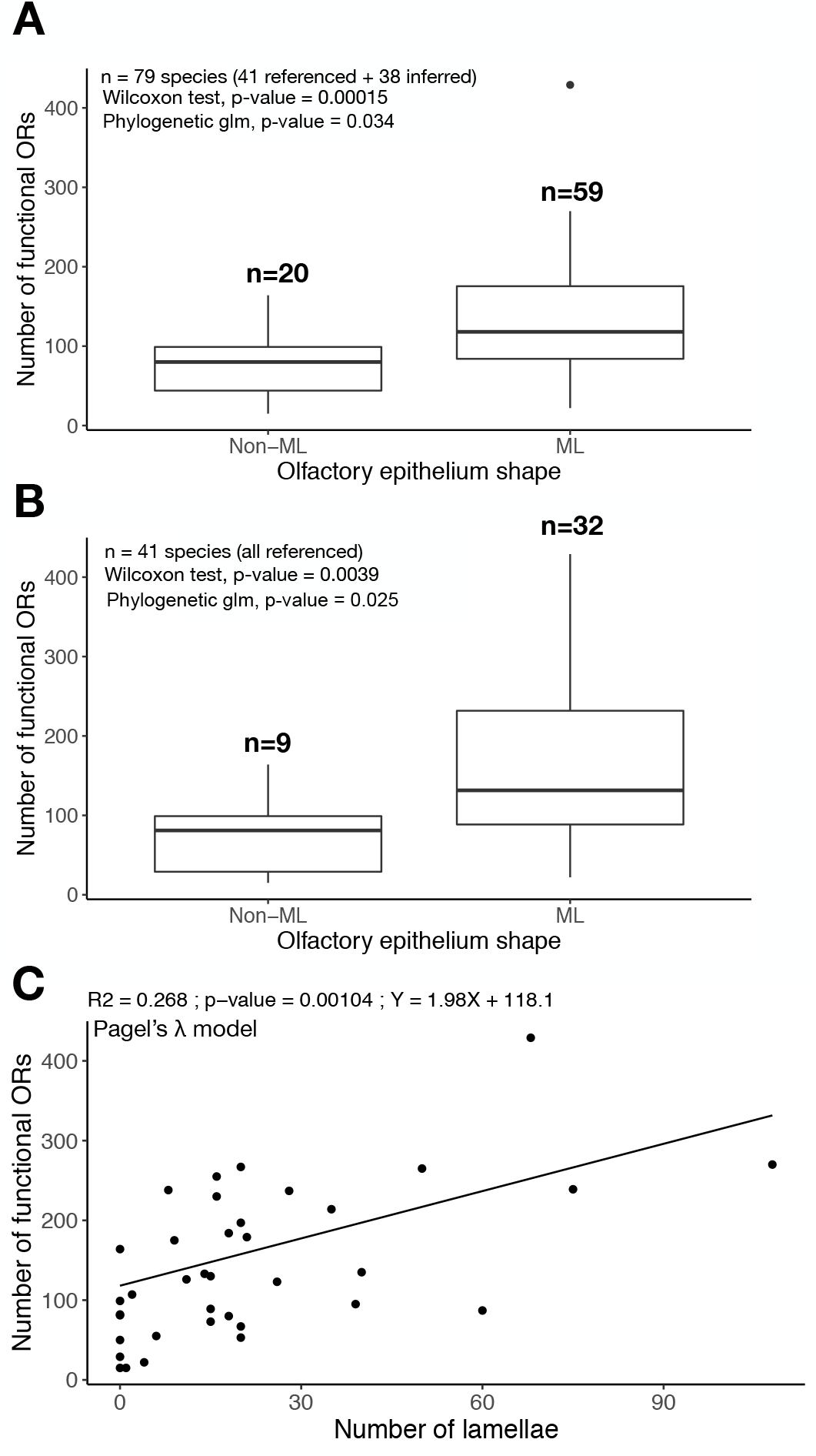
Correlation between the number of functional OR genes and olfactory epithelium characteristics. (A) Correlation with absence (Non-ML) or presence (ML) of a multi-lamellar olfactory epithelium, without taking into account the phylogeny (Wilcoxon test) or taking into account the phylogeny (Phylogenetic glm) and considering species for which data on the olfactory epithelium are either available or are available for a congeneric species. (B) Same analysis as (A), but restricted to species for which data are available. (C) Correlation between the number of lamellae and the number of functional OR genes taking into account the phylogeny with a Pagel’s λ model.

We also found a positive correlation between the number of lamellae and the number of functional OR genes (r = 0.58, p-value = 0.00016), which was also significant after phylogenetic correction with Pagel’s λ model (r = 0.52, p-value = 0.00104, AIC = 432.6) or an Ornstein-Uhlenbeck model (r = 0.49, p-value = 0.002, AIC = 431.2), and a Brownian motion model (r = 0.33, p-value = 0.043, AIC = 435.5) (**fig. 5C**). However, there was no correlation between the number of pseudogenes and OE shape or the number of lamellae (**supplementary fig. S13, Supplementary Material** online).

We tested for a correlation between the number of functional OR genes and other morphological characters (total length of fish, relative eye size), ecological parameters (trophic level, preferred temperature, and maximum depth) which were retrieved from FishBase (https://www.fishbase.us), and with genome size. No significant correlations were found (**supplementary fig. S14, Supplementary Material** online).

## Discussion

We examined the evolutionary dynamics of the size of the OR gene repertoire in a broad sample of teleost fishes and found a larger range of variation than previously recognized (Niimura 2009; Zhu, et al. 2017; Jiang, et al. 2019; Lv, et al. 2019). Large repertoires do not seem to result from bursts of gene duplication, but we identified several episodic high rates of gene pseudogenization that occurred at the root of clades with species that have very few OR genes. Our analysis indicates that the richness of the OR gene repertoire correlates with the presence of a multi-lamellar OE. We found that some large gene losses coincide with a simplification of the OE and that several cases of secondary complexifications of the OE coincide with secondary expansions of the OR gene repertoire.

### OR functions in teleost fishes

In Zebrafish *Danio rerio*, three main types of olfactory sensory neurons (ciliated, microvillus, and crypt) with distinct morphologies, connections and functions, are found in the olfactory epithelium. Segregated neural pathways are responsible for coding and processing information from different types of olfactory stimuli, at least at the level of the olfactory bulb (Friedrich and Korsching 1997). Less is known about molecular components and functional organization of the olfactory system in other teleosts, but similar studies in Channel Catfish *Ictalurus punctatus* (Hansen, et al. 2003) and the Goldfish *Carassius auratus* (Hansen, et al. 2004) indicate that they may be largely conserved in teleosts. OR genes are expressed in ciliated neurons that recognize bile acids and project to medial glomeruli in the olfactory bulb (Yoshihara 2014). Bile acids are biliary steroids synthesized in the liver, secreted in the intestine and released to the environment. These molecules are used as pheromones in the context of social and reproductive behaviors and convey information about the status of the emitter (Doyle and Meeks 2018). For example, one of the most studied and common bile acids, taurocholic acid, acts as an attractant at very low concentration (10^−8^M), to which females of *D. rerio* are more sensitive to than males (Michel and Lubomudrov 1995).

We know virtually nothing about the molecules recognized by the different ORs in teleost fishes because no OR has been deorphanized so far. Although current evidence suggests that OR genes may be expressed in sensory neurons involved in social behaviors (e.g., reproduction, migration, kin recognition, social status) rather than foraging and feeding, more functional data are needed to describe the odorant response range of teleost ORs.

### Diversity and evolution of the OR gene repertoire in teleost fishes

We used the many genomes now available to identify OR gene repertoires in 163 high quality genomes from species representative of teleost diversity. We evaluated the accuracy of our OR gene mining pipeline by comparing our results with similar analyses performed previously on eight species. In comparison to prior studies, we systematically recovered more OR genes, sometimes finding that the initial estimate greatly underestimated the number of OR genes. We found that some species previously thought to have few functional OR genes, such as *Dichotomyctere nigroviridis* (11 genes) and *Ictalurus punctatus* (27 genes) possess larger OR gene repertoires (41 and 87 respectively) and are not among the fishes with the lowest number of OR genes.

We found an unanticipated large range of variation of OR gene number among teleosts, with an average number of 117 OR genes. The lowest number of OR genes (15) was observed in two distantly related species, Broad-nose Pipefish *Syngnathus typhle* and Ocean Sunfish *Mola mola*. In both cases, the few remaining intact ORs primarily belong to the δ and η gene families, which are among the three largest families in teleosts. In both *S. typhle* and *M. mola*, very few pseudogenes were identified, suggesting that contraction of the OR gene repertoire is ancient.

The highest number of OR genes (429) was found in the Zig-zag Eel *Mastacembelus armatus* (Synbranchiformes). This is primarily the result of duplications of genes belonging to δ and ζ families, that are among the three largest OR gene families. The high percentage of pseudogenes indicates high gene turnover in this lineage. Together, these results suggest that in the most extreme cases of reduction and expansion of the OR gene repertoire, the implication of different OR gene families is not strongly biased.

Within teleosts, gene turnover seems to be driven by slow birth-and-death evolutionary dynamics involving similarly low birth and death rates. However, we identified a few instances of high birth and death rates. The largest birth rate was observed at the level of a deep internal branch (**fig. 2**). Because it was followed by various death rates in more recent branches, it is not correlated with extant species that have a rich OR gene repertoire. The second highest birth rate occurred in the common ancestor of *Sander* + *Perca* spp., a split that occurred relatively recently, and, in this case, is associated with high numbers of OR genes, 167 in The Yellow Pech *Perca flavescens* and 217 in the Zander *Sander lucioperca*, compared to the teleost average of 117. More striking is the link between the two highest death rates and the two species with the smallest OR gene repertoires, indicating ancient massive gene losses without compensation by gene duplications in some lineages.

An extensive study of the dynamics of OR gene repertoires in mammals found a low correlation (r = 0.31) between the number of gene losses inferred using the reconciled-tree method and the number of observed pseudogenes (Niimura, et al. 2014). Our analyses suggest an explanation for this counter-intuitive result. Indeed, using all terminal branches, we found a similarly low correlation (r = 0.17) for teleosts. However, using only data for short terminal branches, which corresponds to the most recent evolutionary events in independent lineages, the correlation was much higher (r > 0.7 when using only branches shorter than 7 Myr). This suggests that the oldest pseudogenes are not always identified because they are too divergent or because they are removed from genomes by large deletions, hampering accurate estimation of the number of pseudogenes in long terminal branches. Analyses at smaller phylogenetic scales showed an expected congruence of estimates of the rate of gene losses based on these independent approaches.

### Number of OR genes and correlation with OE complexity

We searched for potential correlations between the size of the OR gene repertoire and morphological traits, ecological parameters and genome sizes. We did not find any correlation with fish length, relative eye size, maximum depth range, trophic level, temperature preference or genome size. In contrast, the shape of the olfactory epithelium (absence or presence of a ML-OE) and, when a ML-OE is present, the number of its lamellae, were positively correlated with the number of OR genes. These correlations suggest that epithelial folding allows for the presence of more sensory neurons but also a more diversified OR repertoire. Moreover, these correlations indicate a relationship between morphological and genomic changes in accordance with the relative importance of olfaction among the suite of sensory systems.

### Case studies

It is instructive to study cases of extreme OR gene repertoire reduction and subsequent evolution at the molecular and morphological levels.

One case concerns the Broad-nosed Pipefish *Syngnathus typhle* and closely related species. The inference of gene gains and losses indicates a large number of gene losses in the common ancestor of the clade containing Syngnathiformes, Kurtiformes and Gobiiformes. If the phylogenetic relationship between Syngnathiformes and Gobiiformes is correct, then the disappearance of the rosette in their common ancestor is closely related with a massive reduction of the OR gene repertoire. In the common ancestor of Syngnathiformes, the number of OR genes also greatly declined. The tiny repertoire of OR genes could be, with a flat olfactory epithelium, a shared character of Syngnathidae (seahorses, pipefishes, and seadragons). Behavioral studies suggest that vision is the leading sense during mating, and that olfaction is less important in *S. typhle* (Berglund and Rosenqvist 2001; Lindqvist, et al. 2011). In Gobiiformes and Kurtiformes, a re-expansion of the OR gene repertoire could have occurred at different rates but no data is available to relate it with a re-acquisition of a multi-lamellar olfactory epithelium.

The diversity of olfactory epithelium within Tetraodontiformes presents a potential model to study evolution of the olfactory system and the OR gene repertoires. The last common ancestor of Tetraodontiformes most likely had a multi-lamellar rosette-shaped olfactory epithelium that is still present in three families, Triacanthodidae, Triacanthidae, and Triodontidae (Tyler 1980; Arcila and Tyler 2017) (see also **supplementary fig. S11 E-F-G-H-I Supplementary Material** online). Repeated losses of the ML-OE occurred within Tetraodontiformes, in particular in Monacanthidae and Tetraodontidae. In both cases, loss of the ML-OE is associated with a contraction of the OR gene repertoire to 45 functional OR genes in the monacanthid *Thamnaconus septentrionalis* and 41 functional OR genes in the tetraodontid *Dichotomyctere nigroviridis. Takifugu* spp., also tetraodontids, have a ML-OE, but the series of folds do not have the ancestral rosette shape, suggesting that it might be a secondary re-acquisition of a ML-OE. We infer an expansion of the OR gene repertoire in the common ancestor of *Takifugu* because the functional OR repertoire size ranges from 79 in *Takifugu flavidus* to 100 in *Takifugu rubripes*. Finally, the small number of functional OR genes (15) in Ocean Sunfish *Mola mola* corresponds with their anatomy. *Mola mola* has minute nostrils almost flush with the surface of the skin (Tyler 1980). We examined an adult specimen of *M. mola* (129 cm TL) and found that the olfactory epithelium is flat, except for one small V-shaped fold on the surface below the incurrent nare (KEB unpublished data). The brain of *M. mola* has greatly reduced olfactory nerves as well as reduced olfactory bulbs (Burr 1928). Together this morphological data and the small number of OR genes suggest that olfaction is particularly regressed in this species.

We observed a 3-fold difference in OR number between the Zebrafish *Danio rerio* and the related cyprinid *Danionella dracula*. The miniaturized and transparent species belonging to *Danionella* have recently attracted the interest of neuroscientists because of the advantages for functional imaging, and of developmental biologists as a model for understanding the regulation of size and developmental heterochronies and truncations (Schulze, et al. 2018; Conway, et al. 2020). *Danionella dracula* and *D. translucida* have small, skull-less brains but rich behavioral repertoires, including acoustic communication. A reduction in social olfactory communication associated with OR gene losses could be related or compensated for by the emergence of a sound-producing apparatus and acoustic communications skills in these species. Such sensory trade-offs could also explain the relatively small repertoire of the electric fishes *Electrophorus electricus* and *Paramormyrops kingsleyae* compared to their closest relatives, because their social interactions may primarily involve electric communication (Gracheva and Bagriantsev 2018). However, additional analyses are required to investigate the importance of sensory trade-offs in the evolution of the size of OR gene repertoires in teleosts.

### Perspectives

What kinds of ecological transitions drove in parallel the loss or re-appearance of a ML-OE and contraction or re-expansion of the OR gene repertoire in different teleost lineages? How can Sea-Horses and Ocean Sunfish thrive with such poor OR repertoires and why do several species of *Takifugu* have a ML-OE and a large OR repertoire relative to other Tetraodontiformes that have a flat OE and a limited OR repertoire? Further studies examining social communication, ecology, life history traits and sensory physiology of these species are needed to answer these fascinating questions.

New high-quality genome sequences are becoming available at an ever-increasing rate, but we found that anatomical data on the olfactory apparatus is rare and that refined descriptions of the olfactory epithelium shape are available only for a handful of model species. Future large scale comparative analyses should include such data for a large sample of teleost species. At the molecular level, an interesting complementary analysis would concern the TAAR gene repertoire. It will be particularly important to examine if the correlation we observed between OR repertoire size and olfactory epithelium shape also holds for the TAAR repertoire size.

## Materials and Methods

### Assessment of genome completeness and OR gene mining

The genomes of 307 teleost species and those of two outgroups (*Acipenser ruthenus* and *Lepisosteus oculatus*) were downloaded from NCBI and ENSEMBL databases on 04/10/2020. The names of the species and the genome assembly versions can be found in **supplementary Data S1, Supplementary Material** online.

For each species, the genome completeness was assessed with BUSCO 3.1.0 (Waterhouse, et al. 2018) using the Actinopterygii odb9 database and the expected genome size extracted from the Animal Genome Size Database (Gregory 2020).

We developed a pipeline (https://github.com/MaximePolicarpo/Olfactory_receptor_genes) to mine olfactory receptor genes in each genome following the procedure of Niimura and Nei (2005) (**supplementary Data S1, Supplementary Material** online). Briefly, tblastn (Ye, et al. 2006) was used with a set of query sequences that are teleost OR genes identified by Y. Niimura (2009) and an e-value threshold of 1e-10 to find genomic regions containing OR genes. Non-overlapping best-hit regions were extracted using samtools (Li, et al. 2009). These sequences were extended 1,000 bp upstream and 1,000 bp downstream and open reading frames (ORFs) larger than 750 bp were extracted using EMBOSS getorf (Rice, et al. 2000). Only ORFs that best matched to a known olfactory receptor against the UniProt database (The UniProt Consortium 2019) were retained. These ORFs were aligned with the teleost OR gene set identified by Niimura (2009) and several outgroup sequences (ora1: A0A0R4IM31.1, taar13: XP_021328050.1, adra2a: NP_919345.2, adrb1: AAI62819.1, galr2b: XP_001339169.1, hrh1: XP_009304002.1 and htr1aa: NP_001116793.1) using MAFFT v7.310 (Katoh, et al. 2005), and a maximum likelihood tree was computed with IQ-TREE (Nguyen, et al. 2015). We only kept genes that clustered with known OR genes by visual inspection of trees and removed 100% identical nucleotide sequences using CD-HIT (Fu, et al. 2012). Retained sequences were classified as functional ORs.

This set of functional ORs was then used for a new round of tblastn with a more stringent e-value of 1e-20. For each genome, best-hit regions were extracted and discarded if overlapping with a functional OR gene region. Otherwise, the best tblastn match was extracted and sequences were classified as follows: 1) « Edge » if the blast target sequence was less than 30 bp away from the scaffold end; 2) « Pseudogene » if the blast target sequence contained at least a stop codon or frameshift; 3) « Truncated » if the sequence was incomplete but without internal stop codons or frameshifts.

Functional, Truncated, and Edge genes were aligned with the teleost OR gene set identified by Niimura (2009) using MAFFT and classified into OR gene families based on their position on a maximum likelihood tree computed with IQ-TREE. Pseudogenes were classified based on their best blastx match.

### Phylogenetic framework and morphological/ecological trait assembly

A time-calibrated phylogeny of ray-finned fishes was downloaded from https://fishtreeoflife.org (Rabosky, et al. 2018) (**supplementary Data S1, Supplementary Material** online). The tree was pruned using the R package « ape » (Paradis and Schliep 2019) to keep only the 165 species whose genomes we used in our study (**supplementary Data S1, Supplementary Material** online). Ecological information on each species was extracted from FishBase (https://www.fishbase.se) using the R package « rfishbase » (**supplementary Data S1, Supplementary Material** online).

We gathered information about the olfactory epithelium (OE) shape (flat epithelium versus multi-lamellar OE) and the number of lamellae per ML-OE in 220 actinopterygians from an extensive literature survey (**supplementary Data S1, Supplementary Material** online). Among these species, genomic data were available for 41 species; for an additional 38 species genomic data were available for at least one species belonging to the same genus. For each species, the olfactory epithelium was classified as multi-lamellar, ML-OE, if the presence of a multi-lamellar OE was described or non-ML-OE if only a few lamellae (1 or 2) or no lamella (flat epithelium) was found. Specimens referenced in the literature were nearly all large juveniles or adults. We did not take into account ontogenetic stage, sexual dimorphism, or individual variation in the olfactory epithelium.

The time-calibrated “fishtreeoflife” was pruned again to retain 205 of the 220 species for which information about the olfactory epithelium was available (15 species were not in the “fishtreeoflife”). The evolution of the OE shape was inferred by maximum likelihood (MPPA with F81 model) and parsimony (with DOWNPASS, DELTRAN or ACCTRAN methods) using PastML (Ishikawa, et al. 2019). Trees were rooted and visualized using iTOL (Letunic and Bork 2007) (**supplementary Data S1, Supplementary Material** online). Species names were validated using Eschmeyer’s Catalog of Fishes (Fricke, et al. 2021).

### Small scale OR genes phylogenies

In order to compare the results of our implementation of a method for identifying OR genes (Niimura 2009) with the results obtained in previous studies, sequences of OR genes from eight teleost species (*Danio rerio, Dichotomyctere nigroviridis, Oryzias latipes, Gasterosteus aculeatus, Takifugu rubripes, Siniperca chuatsi, Pseudoliparis swirei* and *Ictalurus punctatus*) were retrieved from these studies (Niimura 2009; Gao, et al. 2017; Jiang, et al. 2019; Lv, et al. 2019). These sequences were aligned with those found in the present study using MAFFT (on amino acids) and a ML tree was computed with IQ-TREE. The optimal model was inferred with ModelFinder, while the robustness of the nodes was evaluated with 1,000 ultrafast bootstraps. Trees were visualized, rooted and annotated using iTOL.

In order to obtain an overall picture of the diversity of OR gene families in teleosts, we also inferred a phylogenetic tree of all functional OR sequences from 38 species, each belonging to a different order (species and order names can be found in **supplementary Data S1, Supplementary Material** online). Protein sequences were aligned using MAFFT and a near-ML tree was computed with FastTree (Price, et al. 2010) and the robustness of the nodes was evaluated with 1,000 fast-global bootstraps.

### OR birth and death rates

We inferred the number of gene gains and losses along each branch of the species tree using a gene tree - species tree reconciliation method. For each OR gene family, an alignment was performed with MAFFT and a maximum likelihood tree was computed with IQ-TREE. Nodes with low bootstrap values (<90%) were collapsed into polytomies using the R package « ape ». We then used NOTUNG 2.9.1.5 (Chen, et al. 2000) to root and reconcile gene trees with the species tree using the phylogenomics option.

Birth and death rates along each branch as well as the mean birth and death rate were computed using the equations of Niimura et al. (2014). We excluded branches with lengths < 2 million years because differences in gene retrieval and genome qualities had a major impact on inferred birth and death rates (**supplementary Data S1, Supplementary Material** online).

### Phylogenetic Generalized Least-Squares analyses

The phylogenetic signal of each trait was estimated with the function phylosig in the R package « phytools » with the option « test = TRUE » (Revell 2012).

The R package « phylolm » (Tung Ho and Ané 2014) was used to perform phylogenetic linear regressions between continuous traits using the phylolm function with various evolutionary models, and phylogenetic generalized linear models between binary and continuous traits using the phyloglm function.

### Correlation between the number of gene losses and the number of pseudogenes in terminal branches

For each terminal branch of the species tree, the number of gene losses was inferred using the gene family trees of intact OR genes and the program NOTUNG 2.9.1.5 (Chen, et al. 2000) as described above. We also estimated the numbers of OR pseudogenes (OR sequences with at least one LoF mutations), truncated genes (most likely pseudogenes resulting from large genomic deletions) and edge genes (truncated genes most likely resulting from assembly artifacts).

The correlation (Pearson’s r) between the number of gene losses and the number of pseudogenes and truncated genes, together or separately, and the number of edge genes was computed for all terminal branches or using only a subset of branches shorter than a given threshold ranging from 2 Myr to 200 Myr.

In order to estimate the time necessary for making a pseudogene undetectable through the accumulation of substitutions and small indels, we ran simulations in which an intact OR sequence accumulated nucleotide substitutions (at a rate of 10-8 per generation) and indels at a rate of 0.05 × 10^−8^per generation as estimated previously (Policarpo, et al. 2020). We also took into account the transition/transversion ratio and the relative frequency of indels ranging from 1 and 9 bp in length (more details on the numerical values used for the simulations are given in the Python script available at https://github.com/MaximePolicarpo/Olfactory_receptor_genes).

Each time a new mutation was produced in the evolving sequence, a tblastn analysis was performed using the evolving DNA sequence and the translated original OR sequence. The evolving sequence was classified as « unrecognized » if the e-value was greater than 10^−20^, which was the threshold used to mine OR pseudogenes, and the number of generations to get an unrecognized sequence registered. Running this simulation 10,000 times allowed the estimate of the distribution of the number of generations necessary to make a pseudogene undetectable.

### Data availability

All the data and the results of the analyses performed in this study are available for download at figshare (https://figshare.com/articles/dataset/Supplementary_files_-_ORs_in_teleost/14180090): Nucleotide sequences of the olfactory receptor genes in three FASTA format files (functional genes, incomplete genes and pseudogenes); Version of genomes used for this study, assembly sizes, expected genome sizes, BUSCO assessment results, olfactory receptor gene count, olfactory epithelium shape, ecological data as well as NOTUNG results in Supplementary_Table.xlsx; BUSCO assessment results in graphic format and phylogenetic trees (Newick format).

## Supplementary Material

Supplementary data are available at Molecular Biology and Evolution.

## Acknowledgments

We thank Matthieu Haudiquet for his help with the R package « phylolm »

## Notes

### Competing Interest Statement

The authors have declared no competing interest.

