## Supplementary fig. S1 for "Evolutionary dynamics of the OR gene repertoire in teleost fishes: evidence of an association with changes in olfactory epithelium shape"

**A**

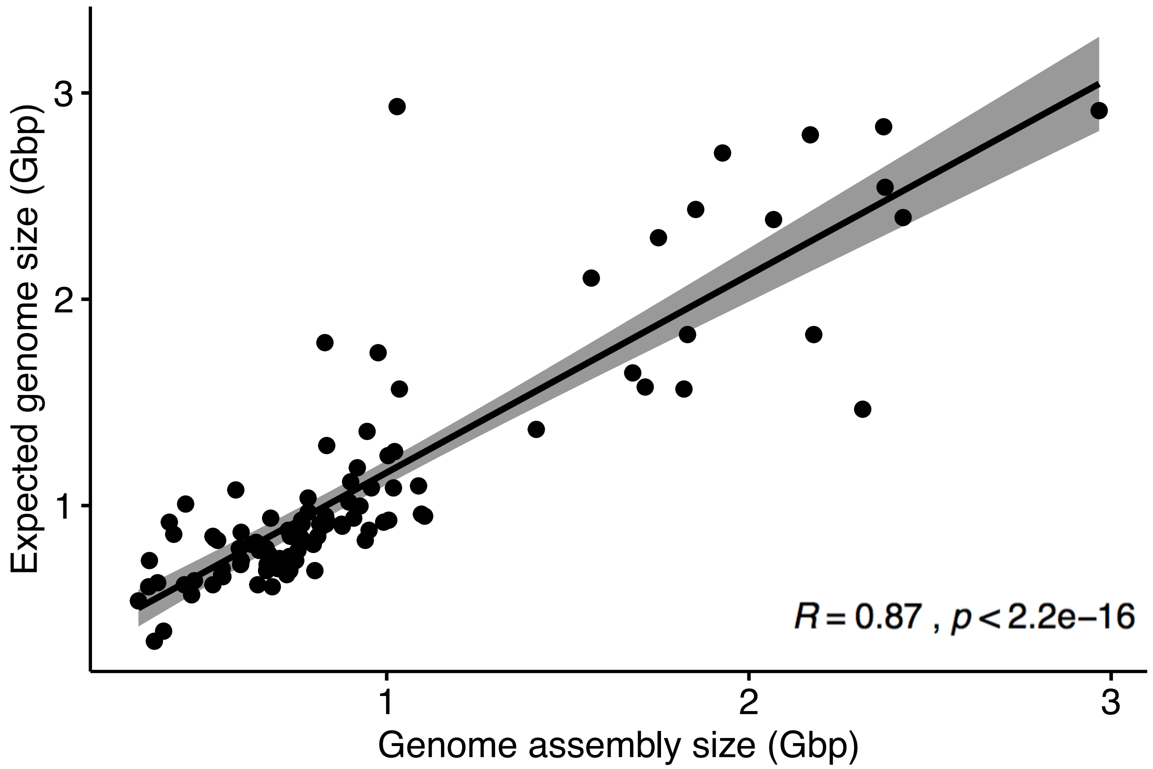


**B**

| Species | Assembly size (bp) | Expected size (bp) | Difference (bp) |
|---|---|---|---|
| *Chionodraco hamatus* | 829,466,164 | 1,789,740,000 | -960,273,836 |
| *Colossoma macropomum* | 2,314,107,020 | 1,467,000,000 | 847,107,020 |
| *Nerophis ophidion* | 976,361,201 | 1,740,840,000 | -764,478,799 |
| *Oncorhynchus nerka* | 1,927,125,257 | 2,709,060,000 | -781,934,743 |
| *Opsanus beta* | 1,028,783,780 | 2,934,000,000 | -1,905,216,220 |
| *Salvelinus alpinus* | 2,169,536,488 | 2,797,080,000 | -627,543,512 |
