## Supplementary figures and images for "Evolutionary dynamics of the OR gene repertoire in teleost fishes: evidence of an association with changes in olfactory epithelium shape"

### Supplementary fig. S2

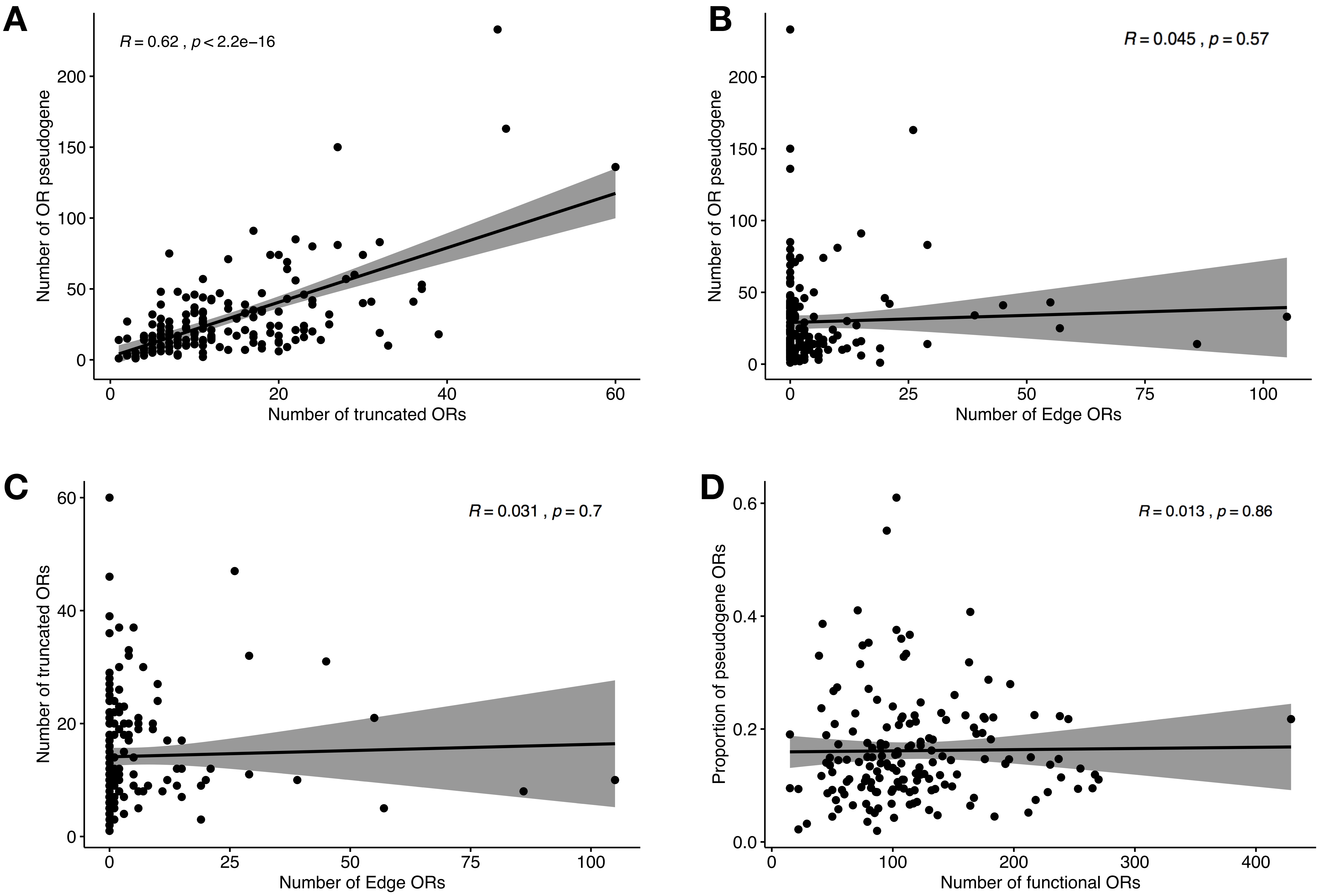

### Supplementary fig. S3

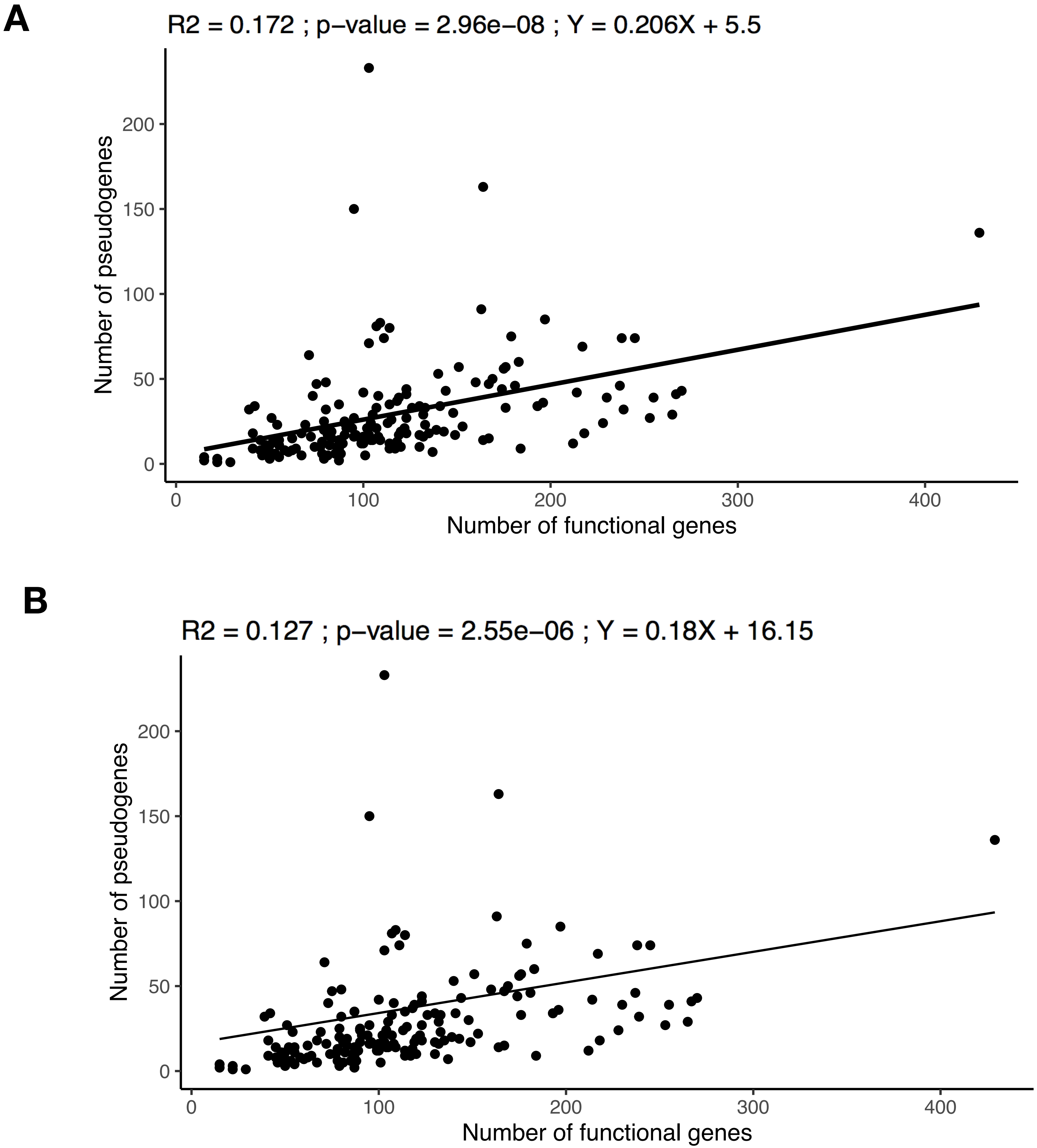

### Supplementary fig. S4

Fig. S4

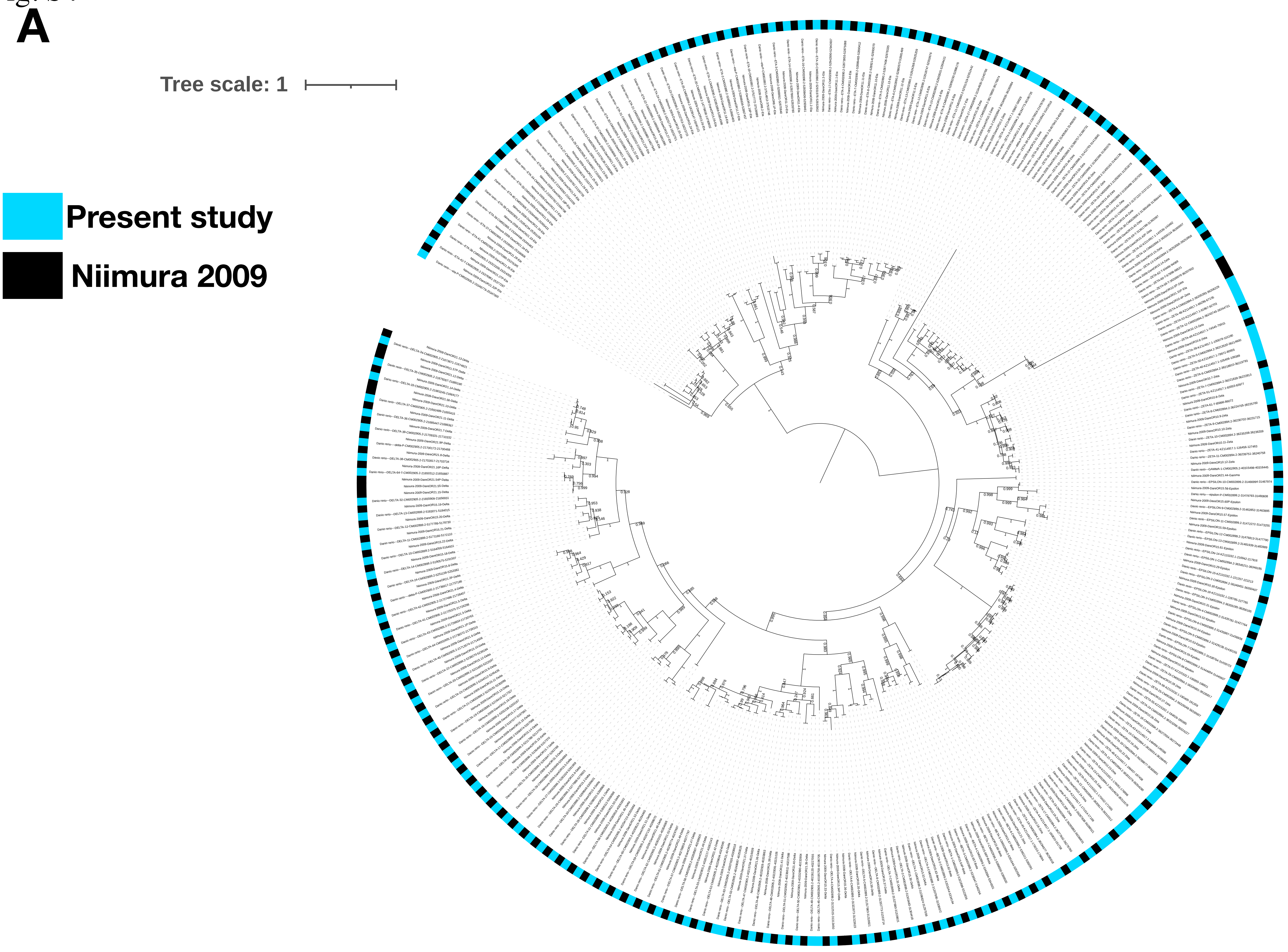

B

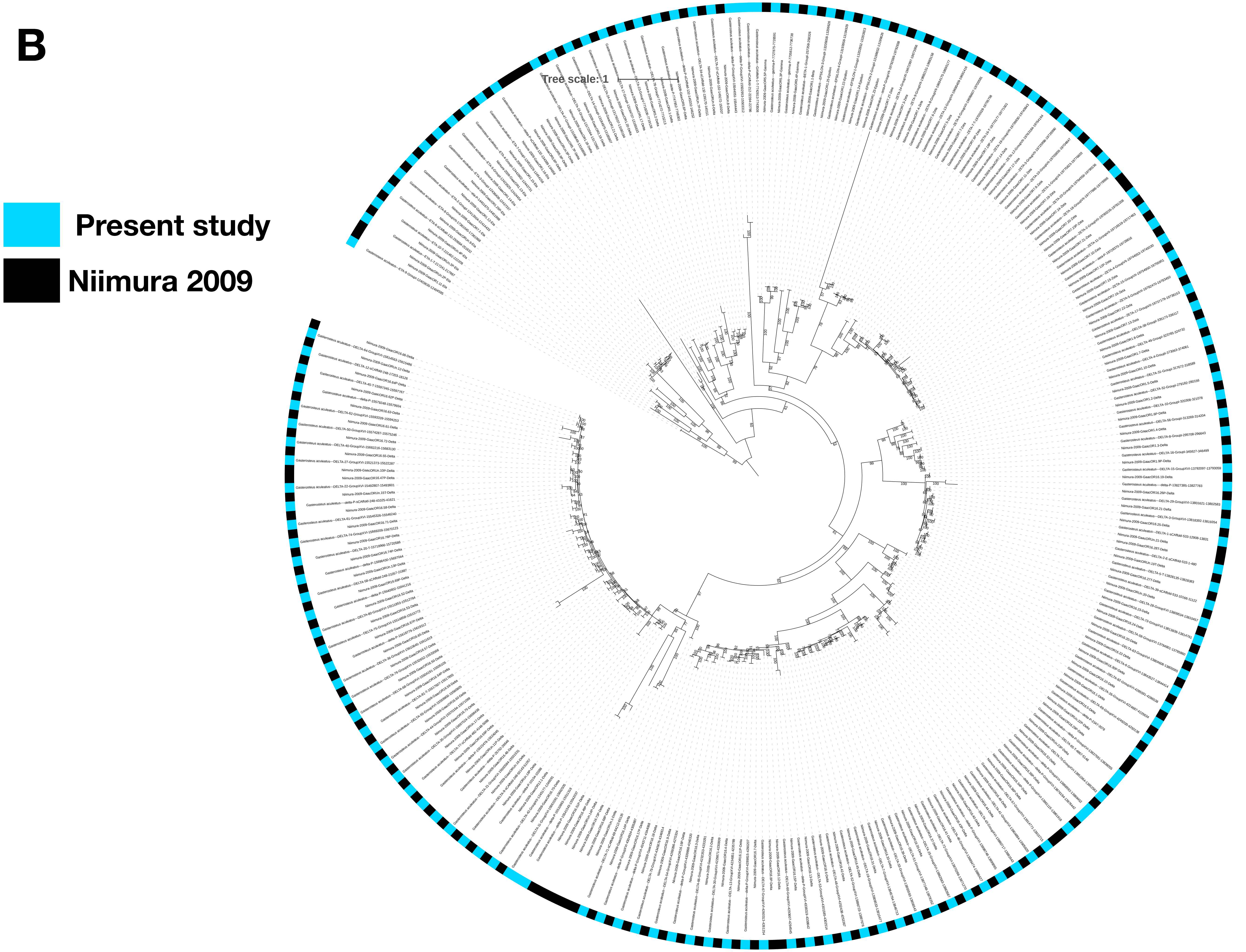

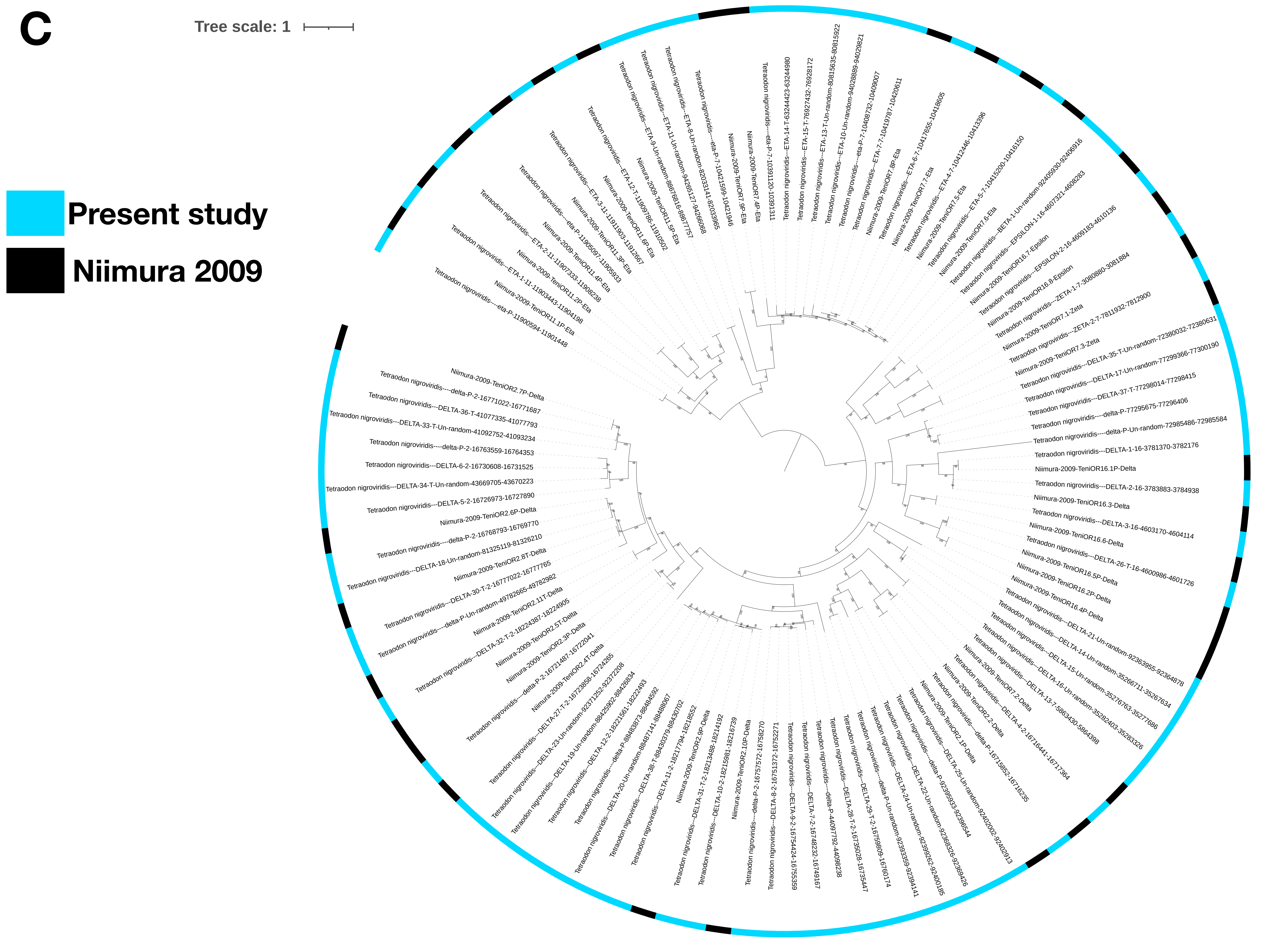

D

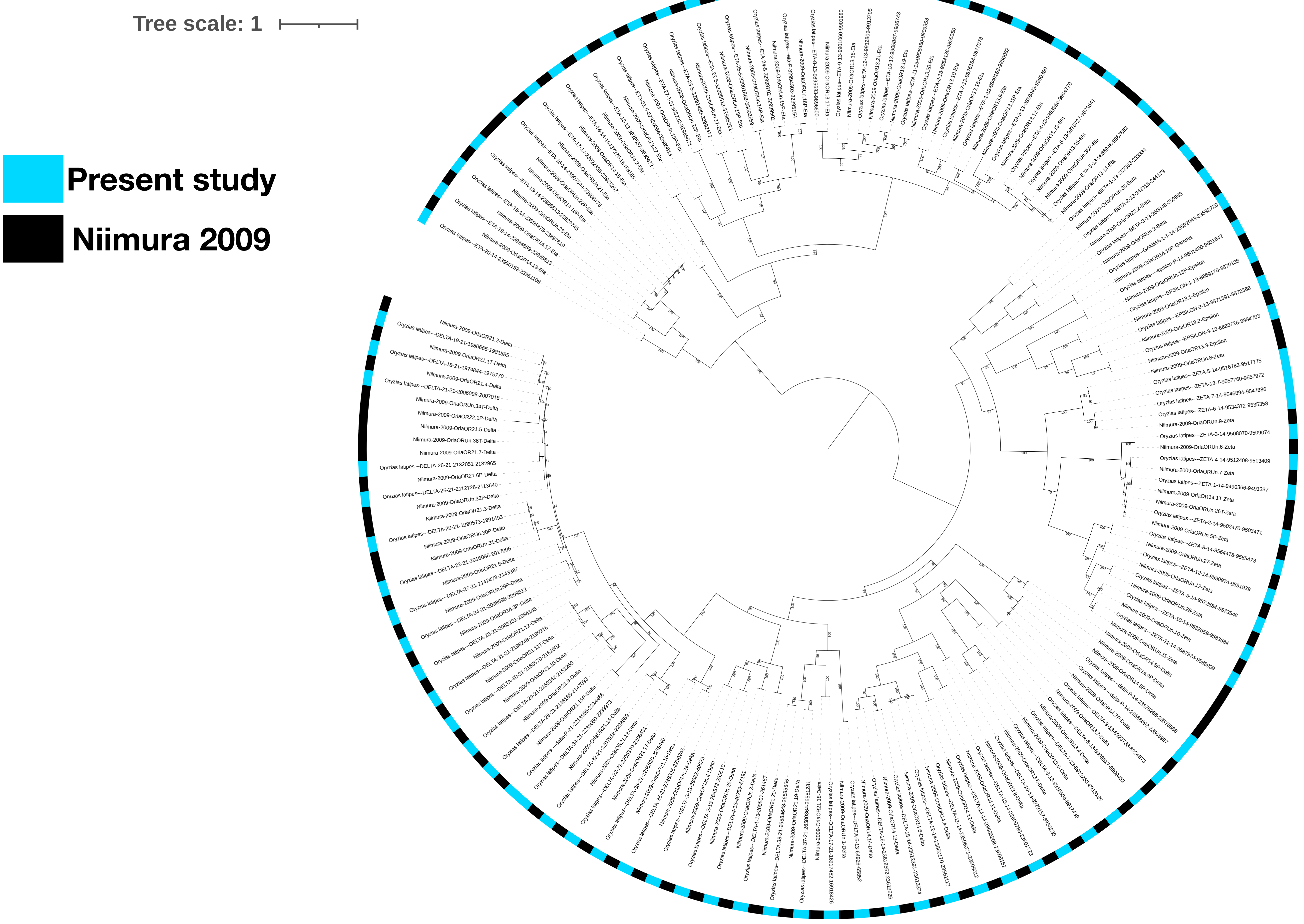

**E**

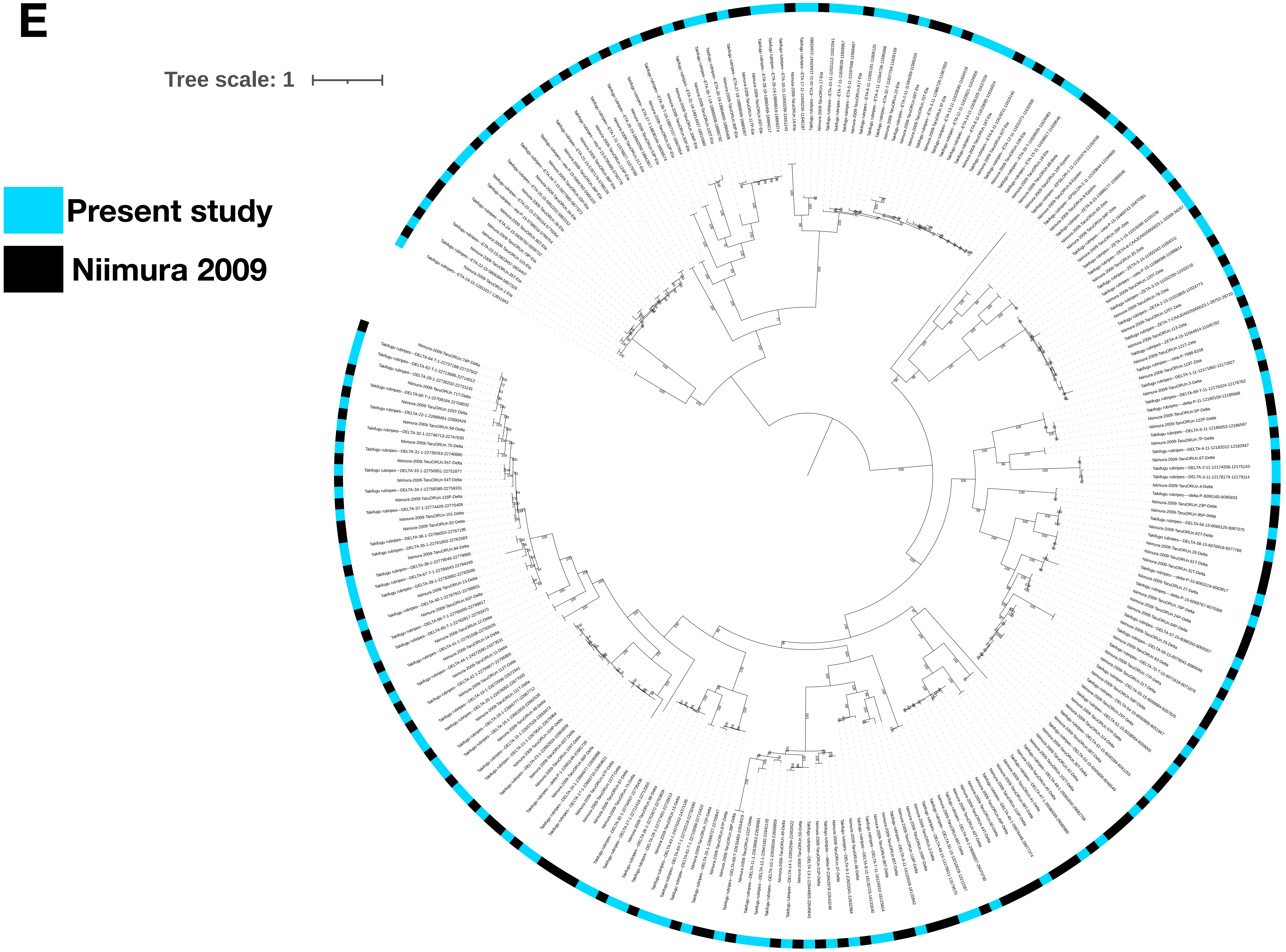

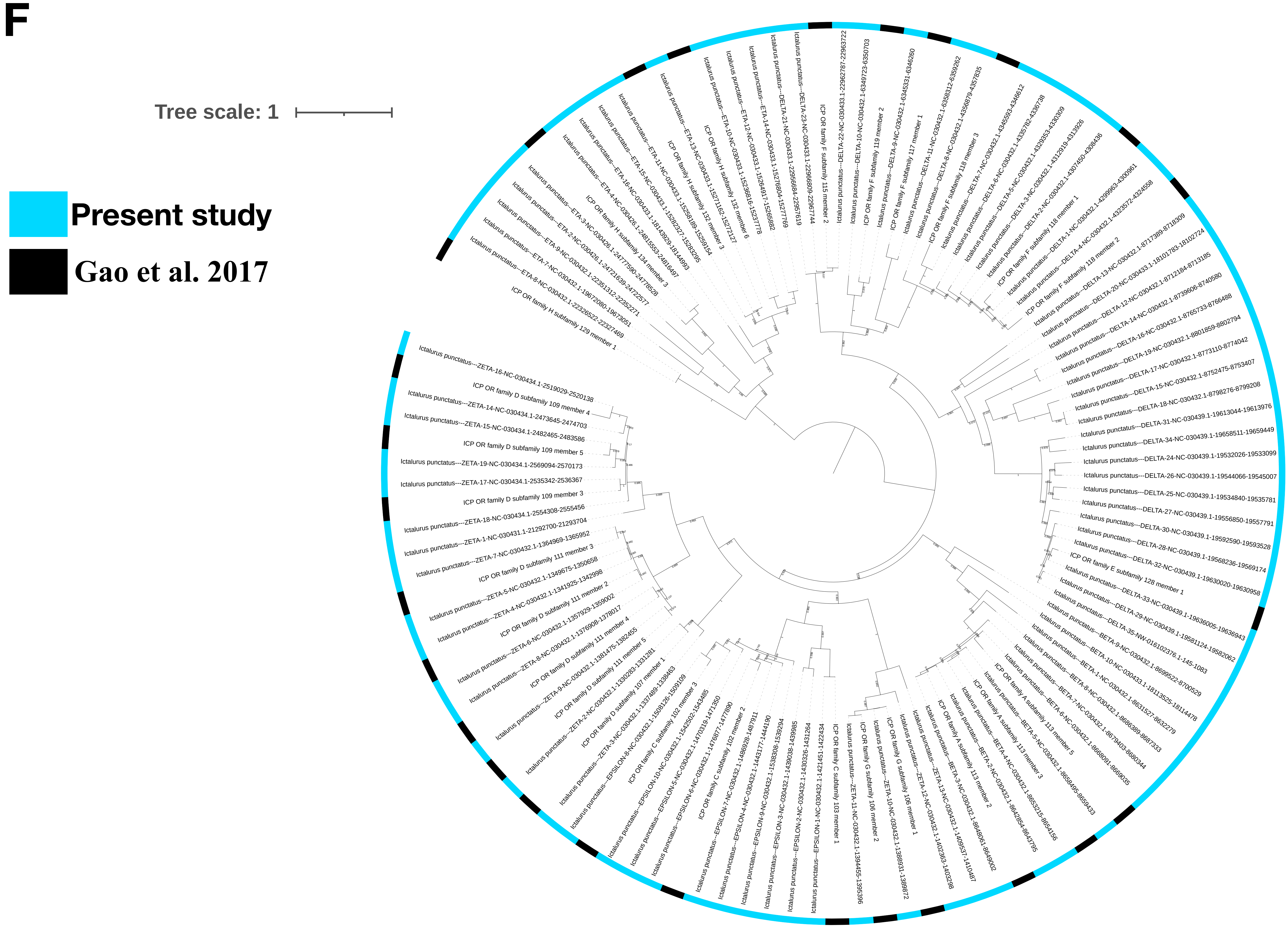

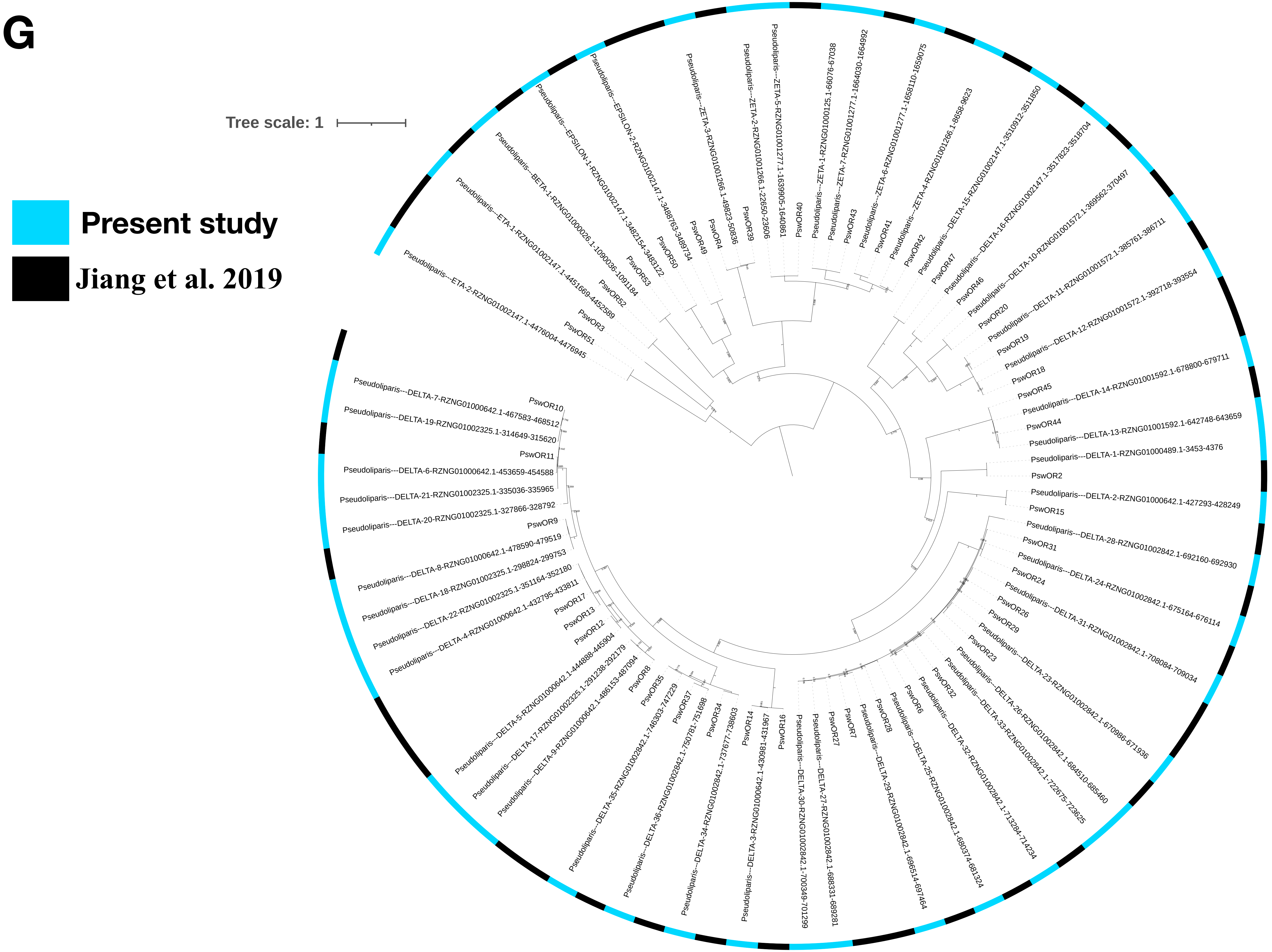

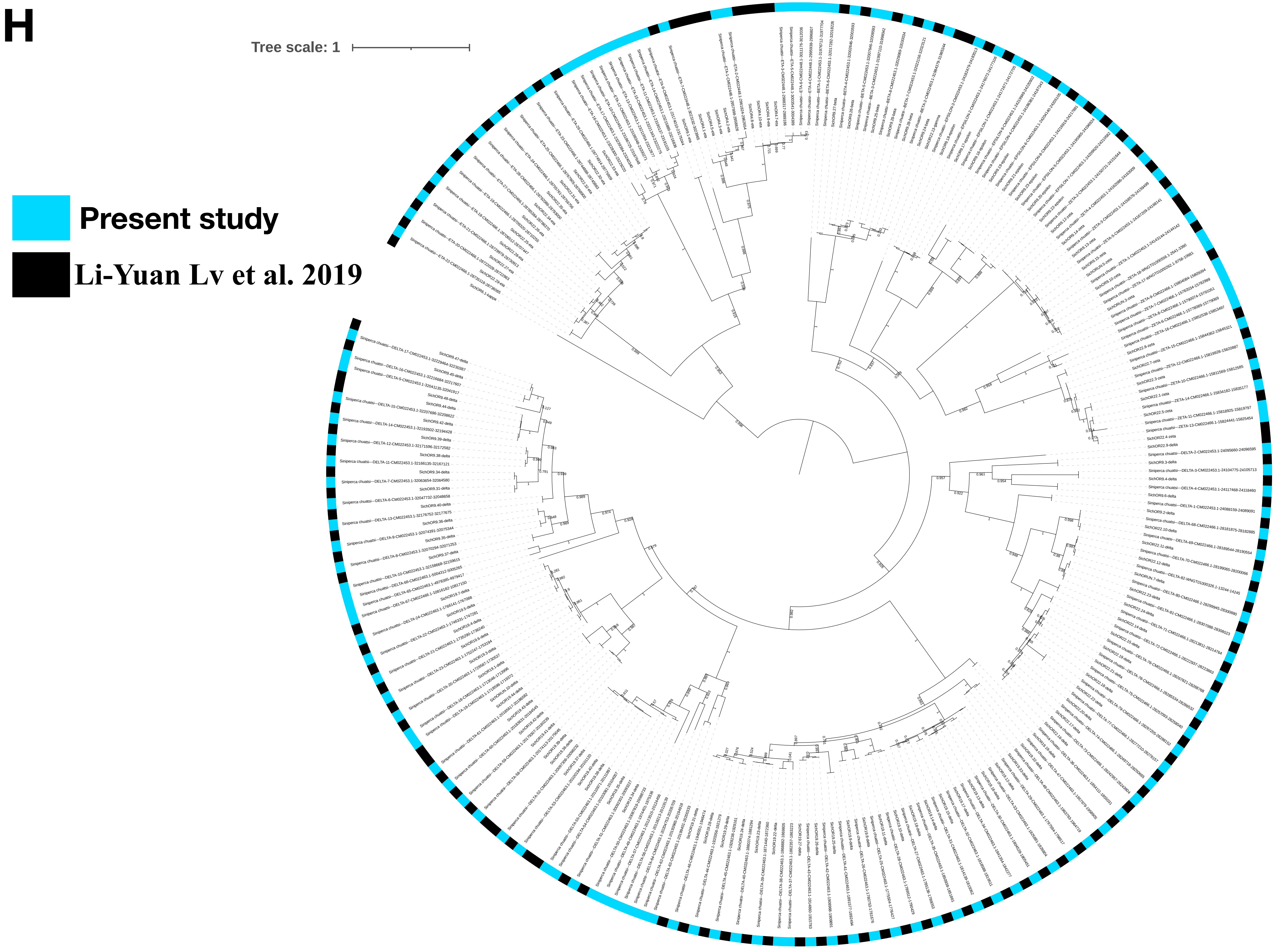

### Supplementary fig. S5

Fig. S5

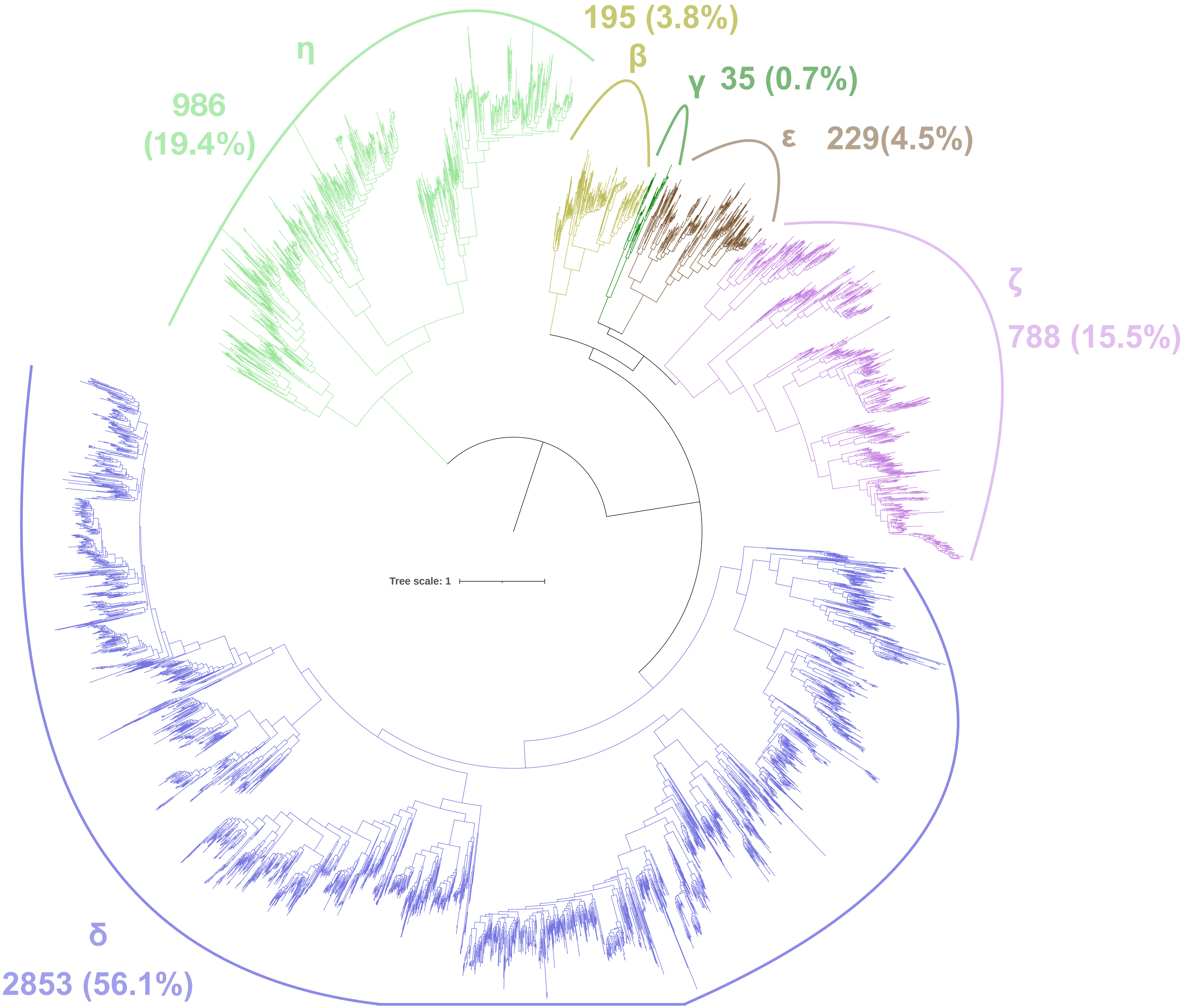

### Supplementary fig. S6

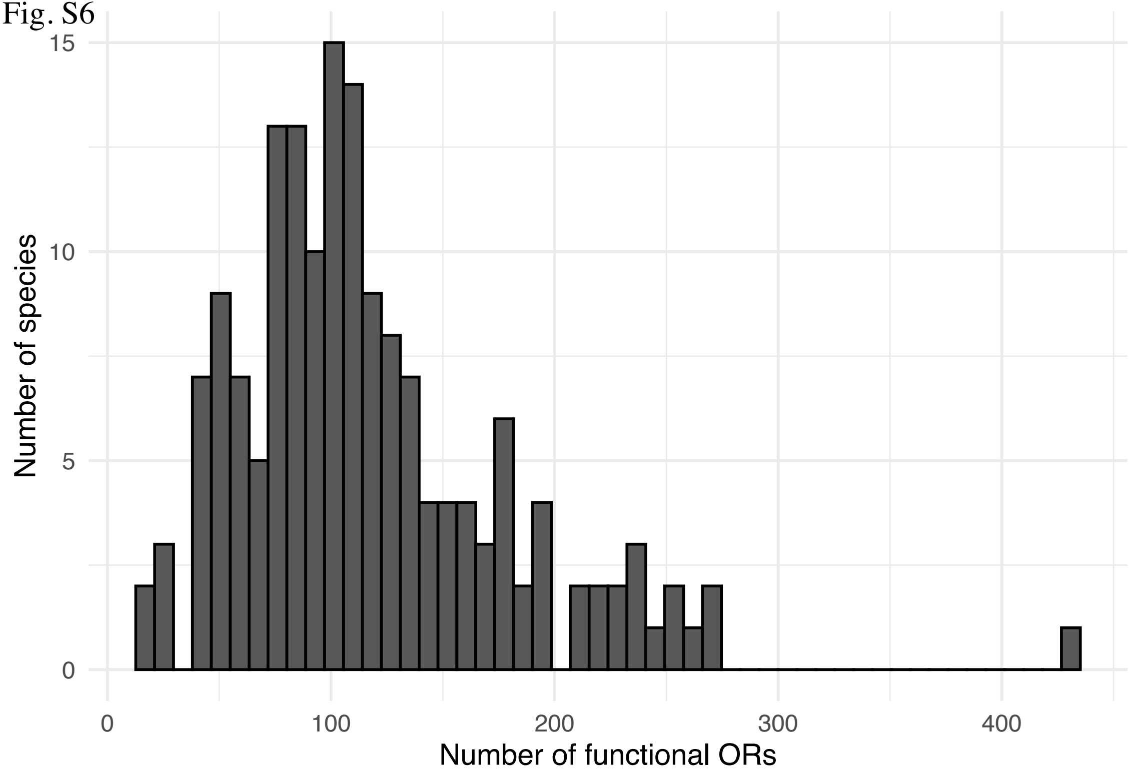

### Supplementary fig. S7

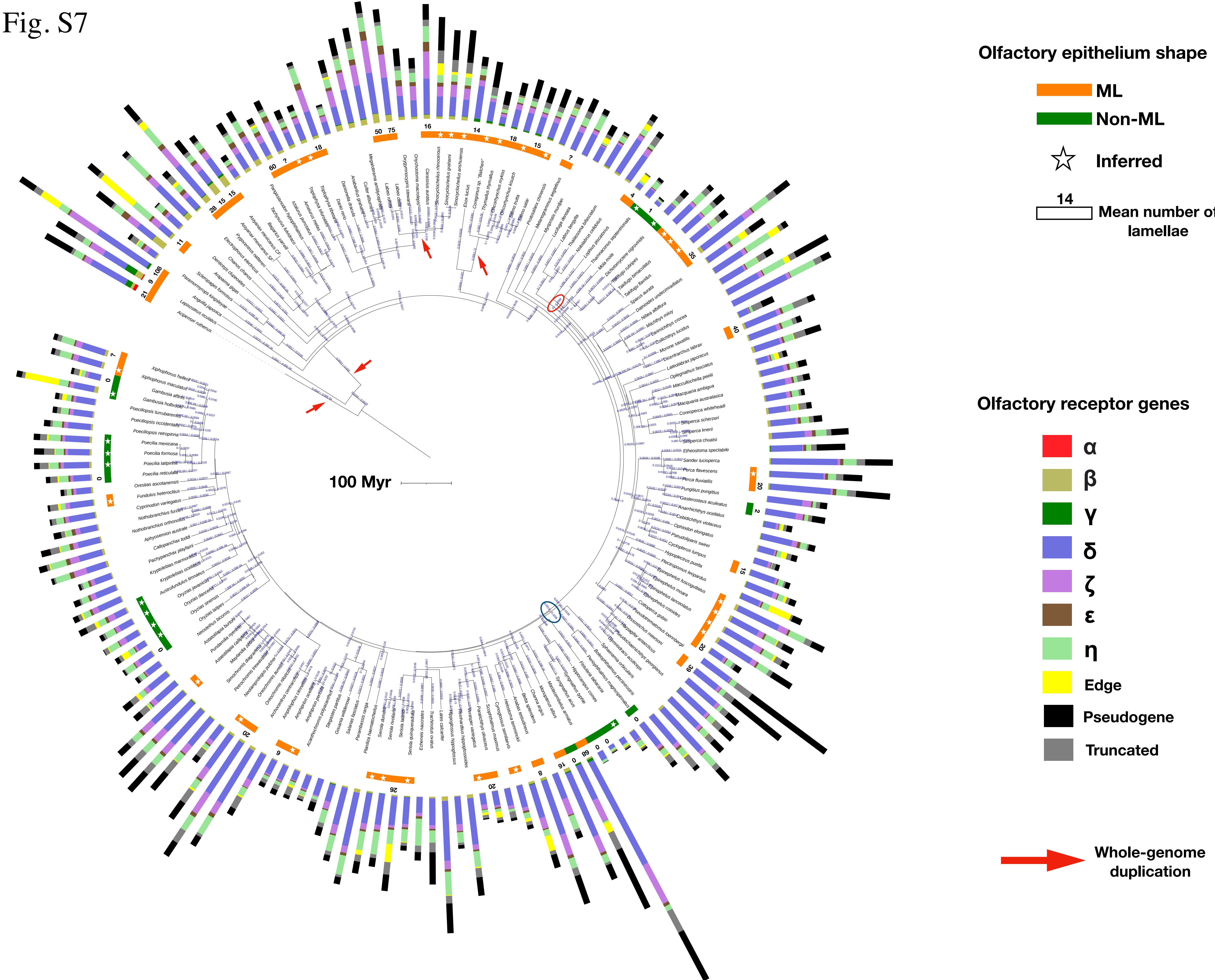

### Supplementary fig. S8

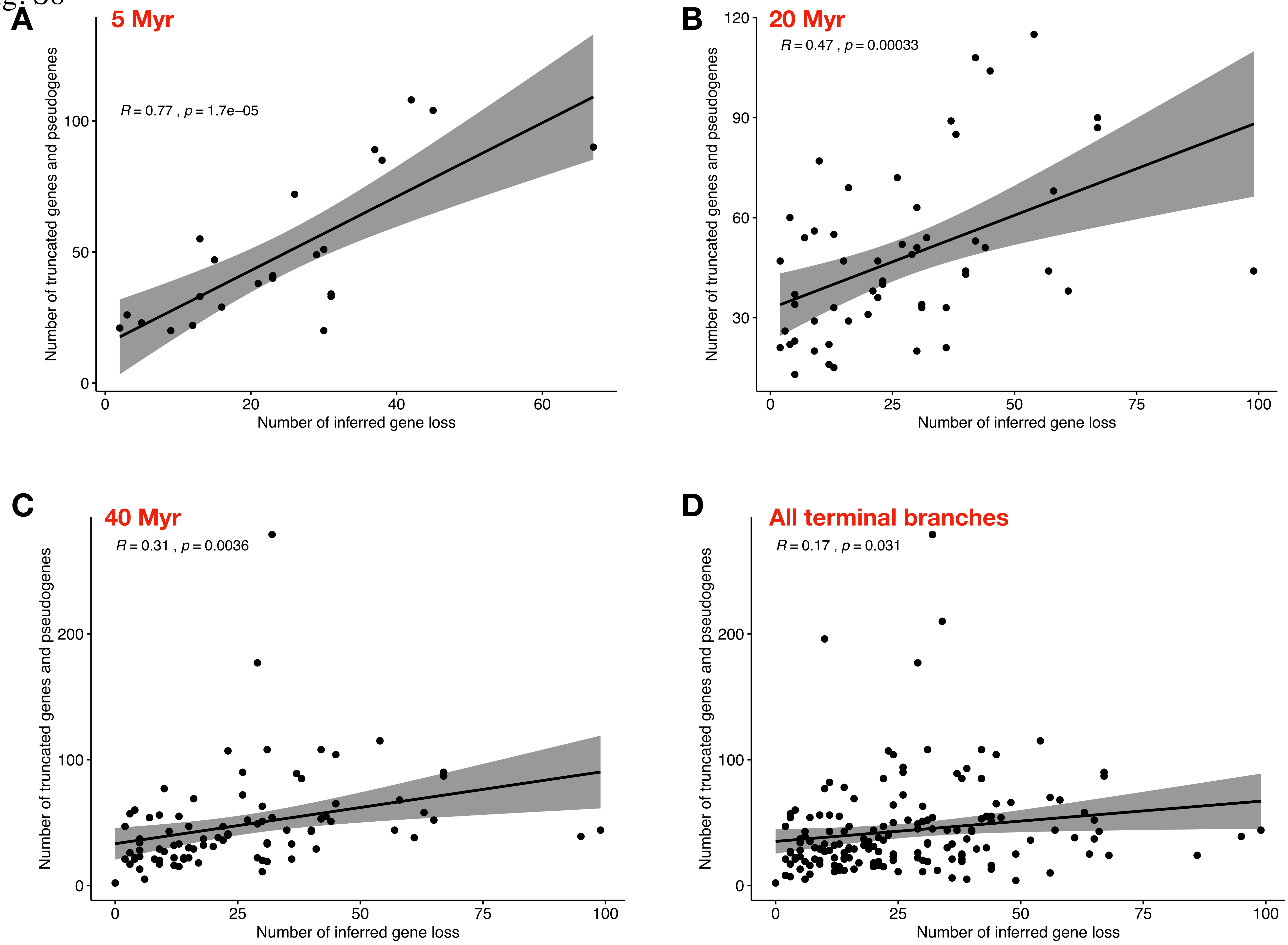

### Supplementary fig. S9

Fig. S9

##### A- Pseudogenes

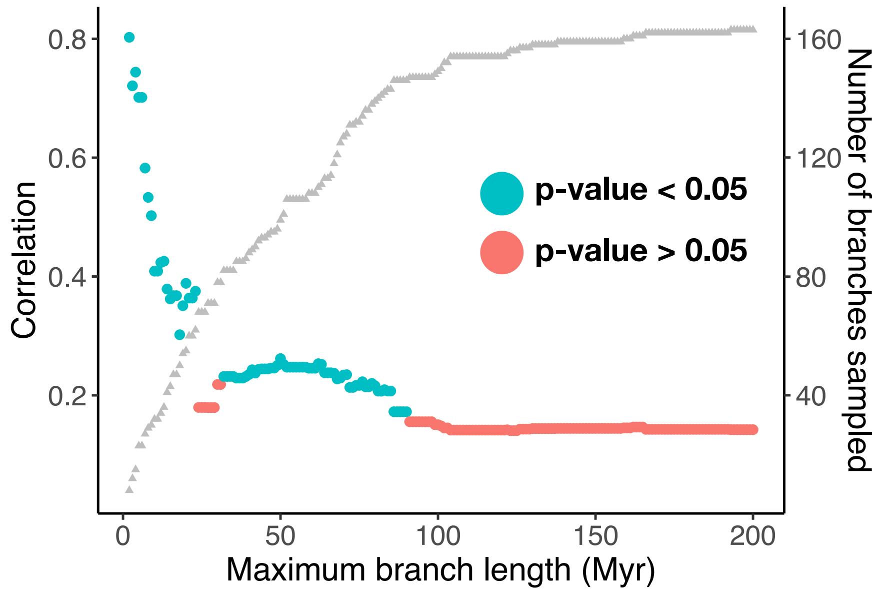


##### B- Truncated genes

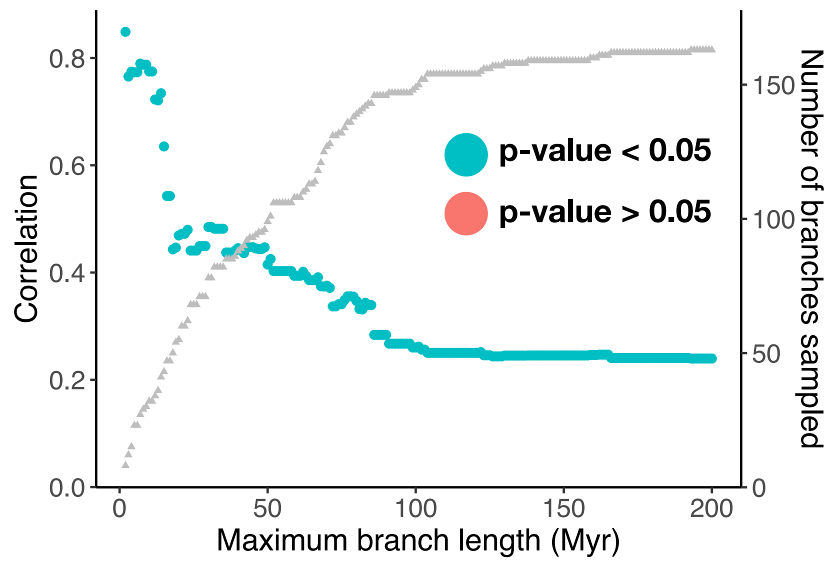


##### C- Edge genes

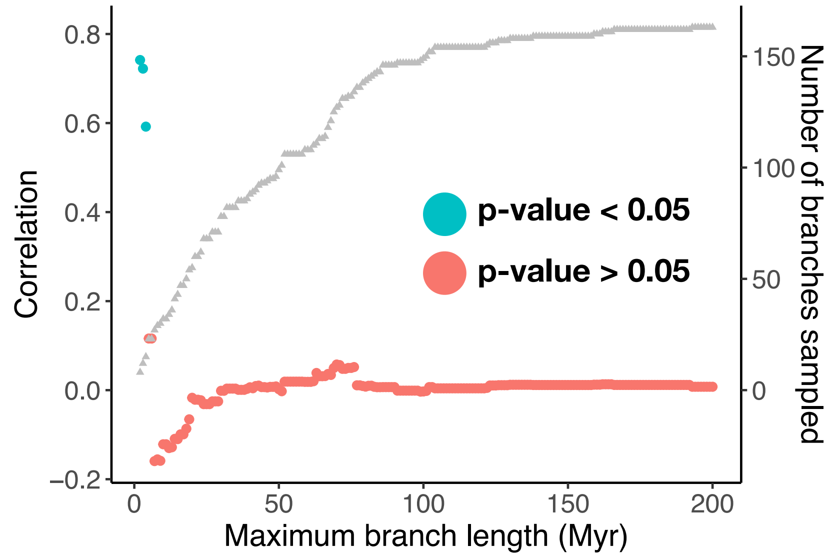

### Supplementary fig. S10

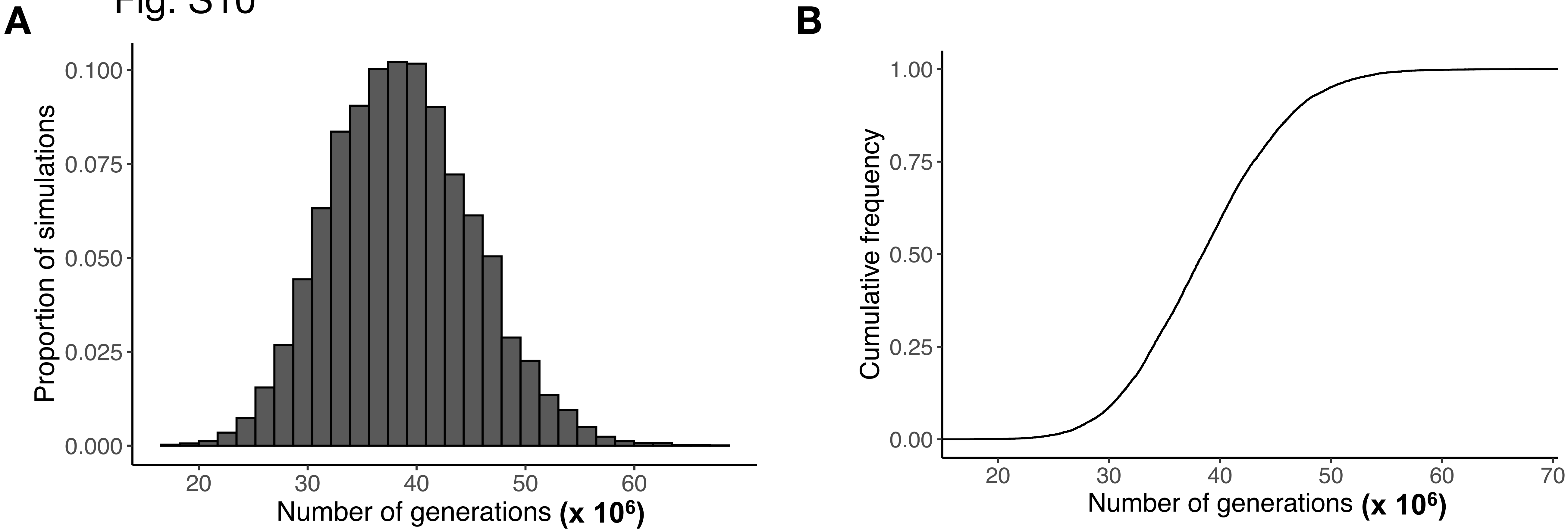

### Supplementary fig. S11

**A - MPPA + F81**

Fig. S11

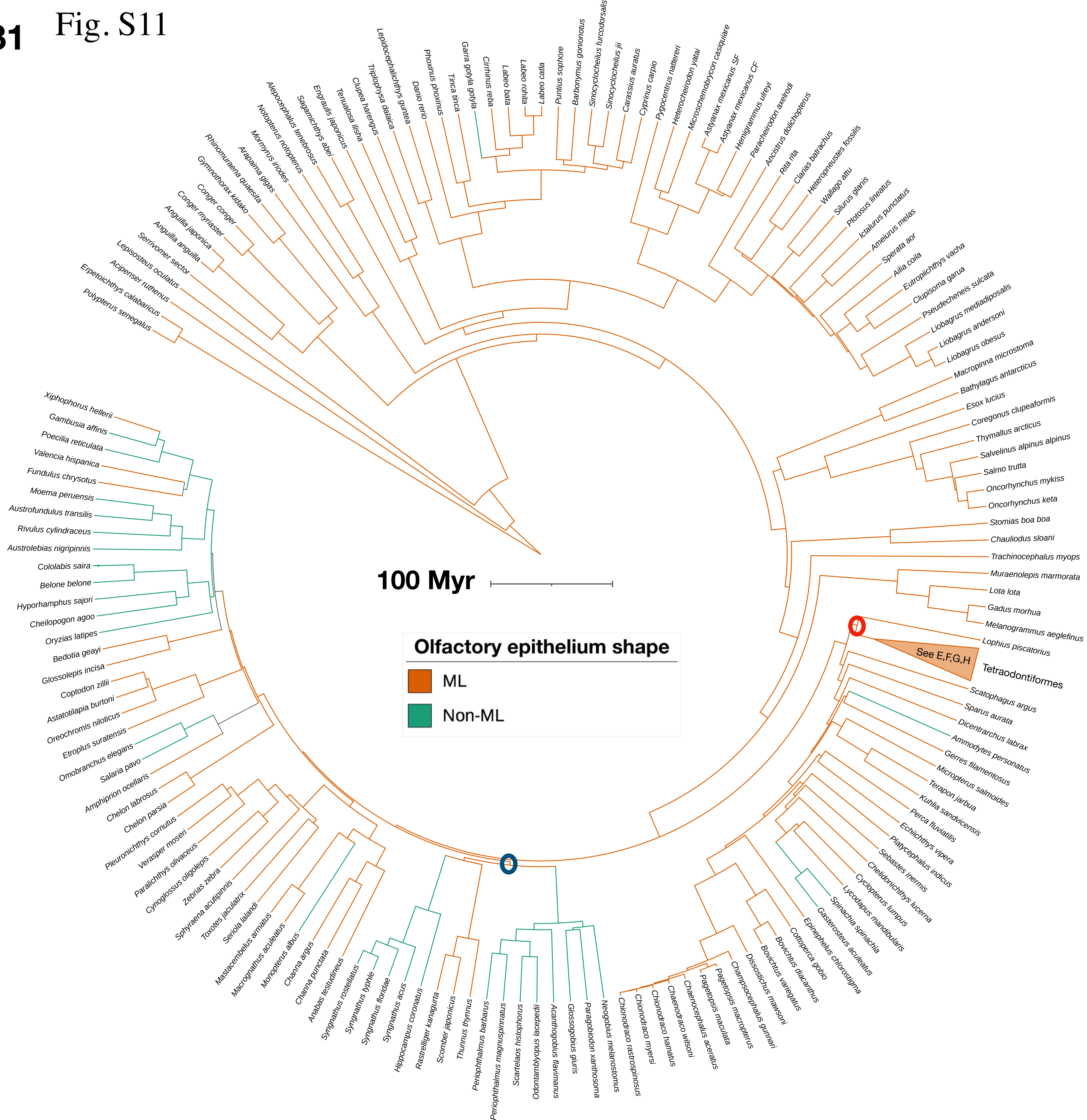

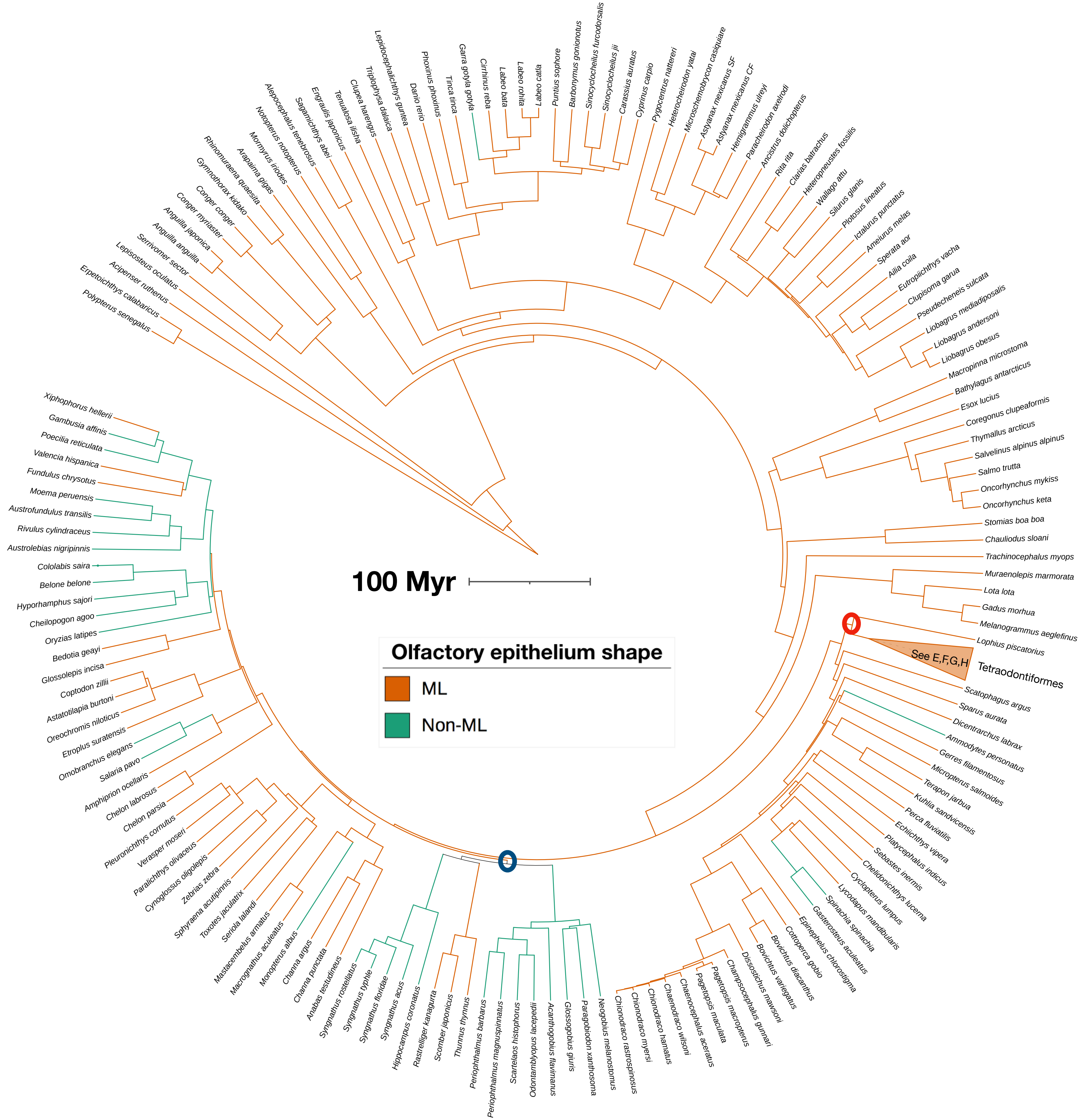

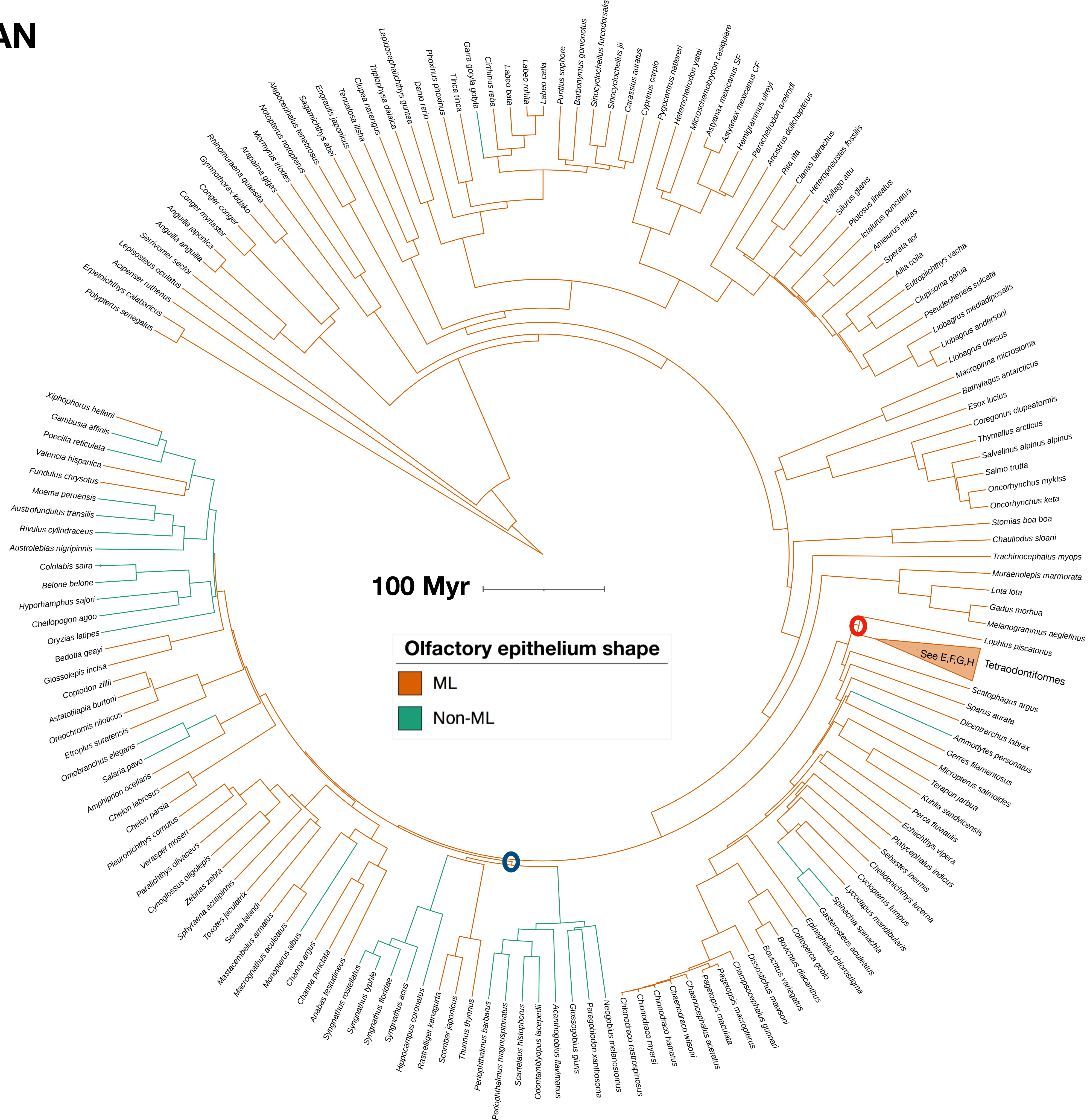

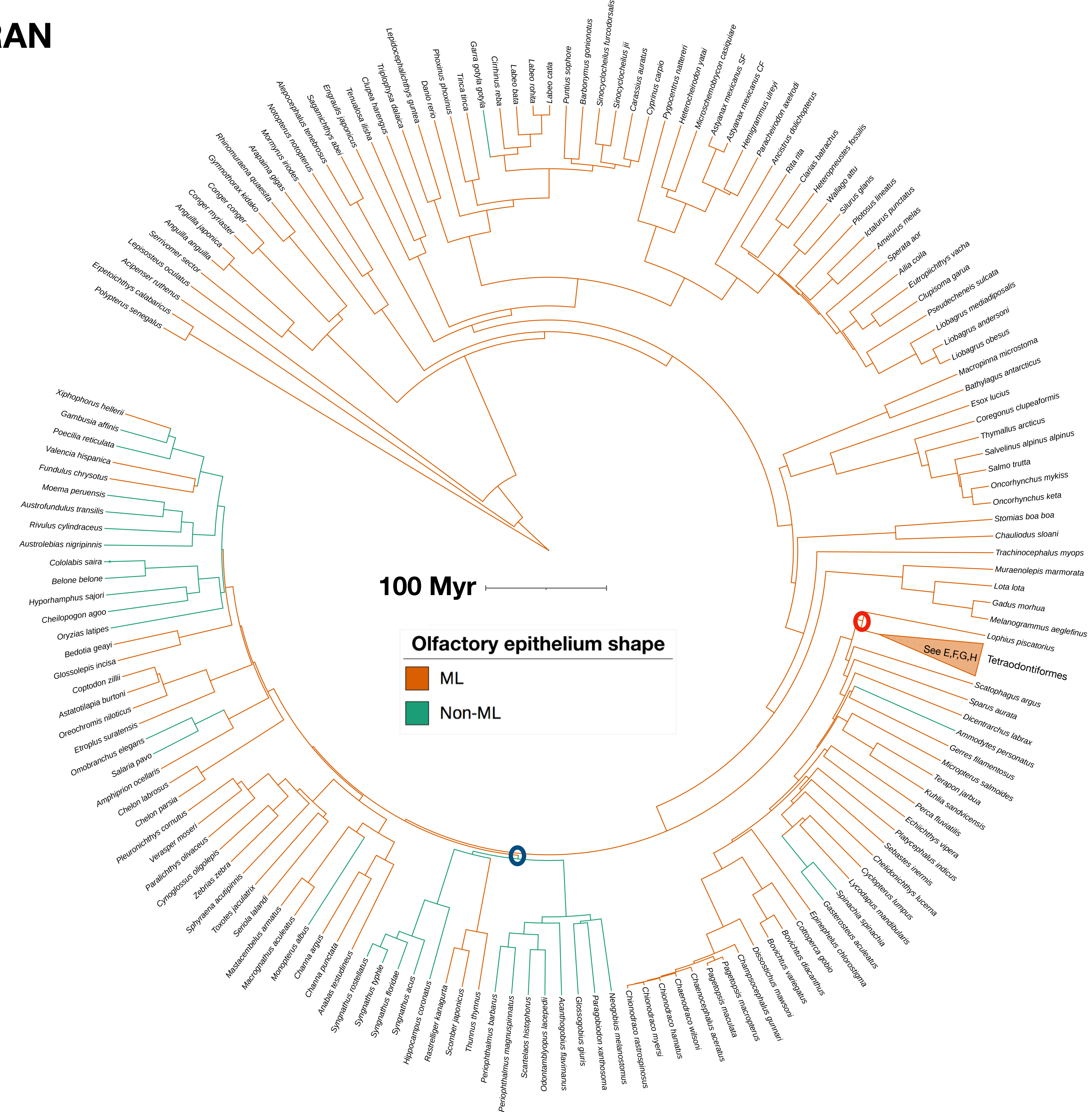

#### E - MPPA + F81 , DOWNPASS, DELTRAN, ACCTRAN

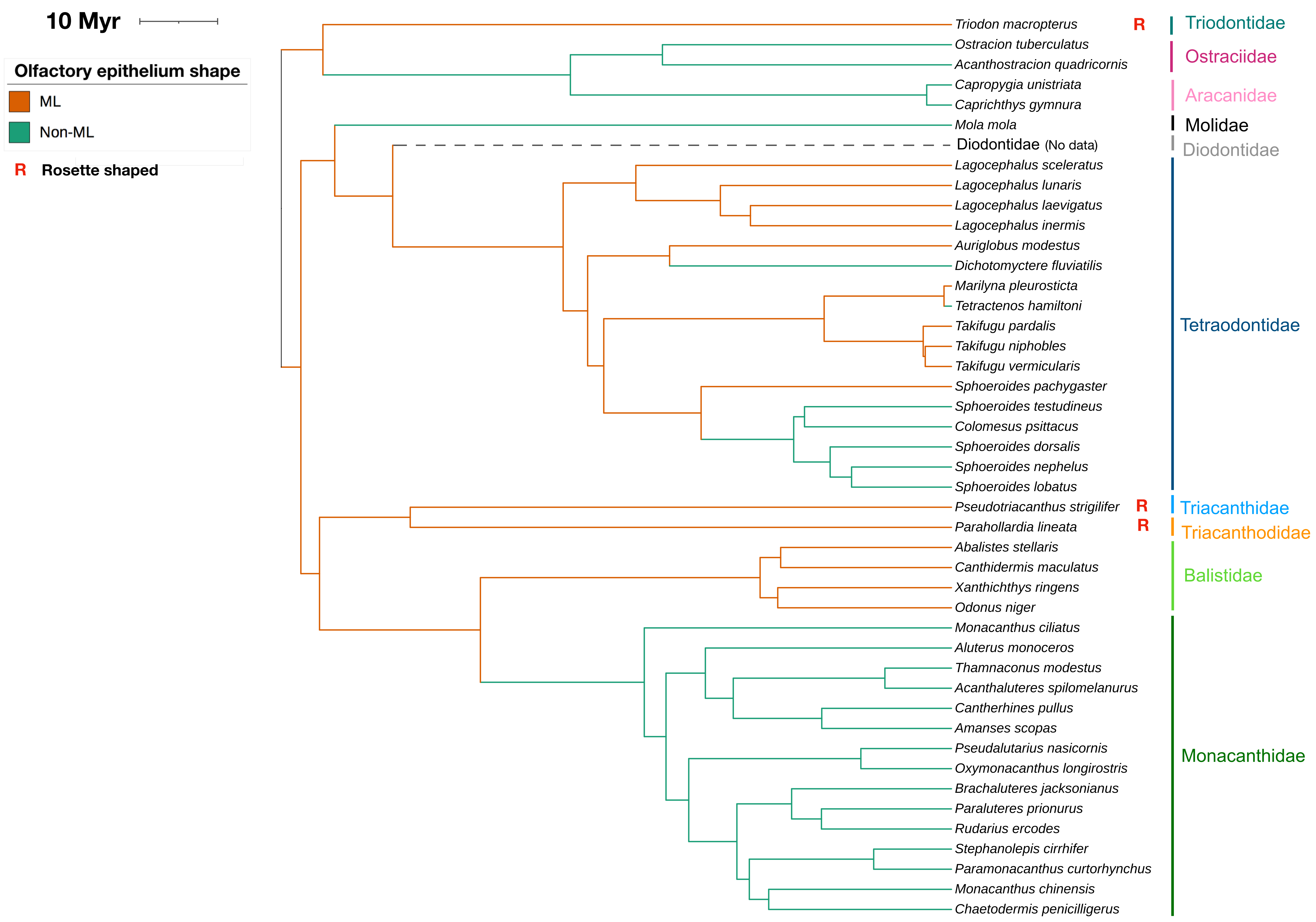

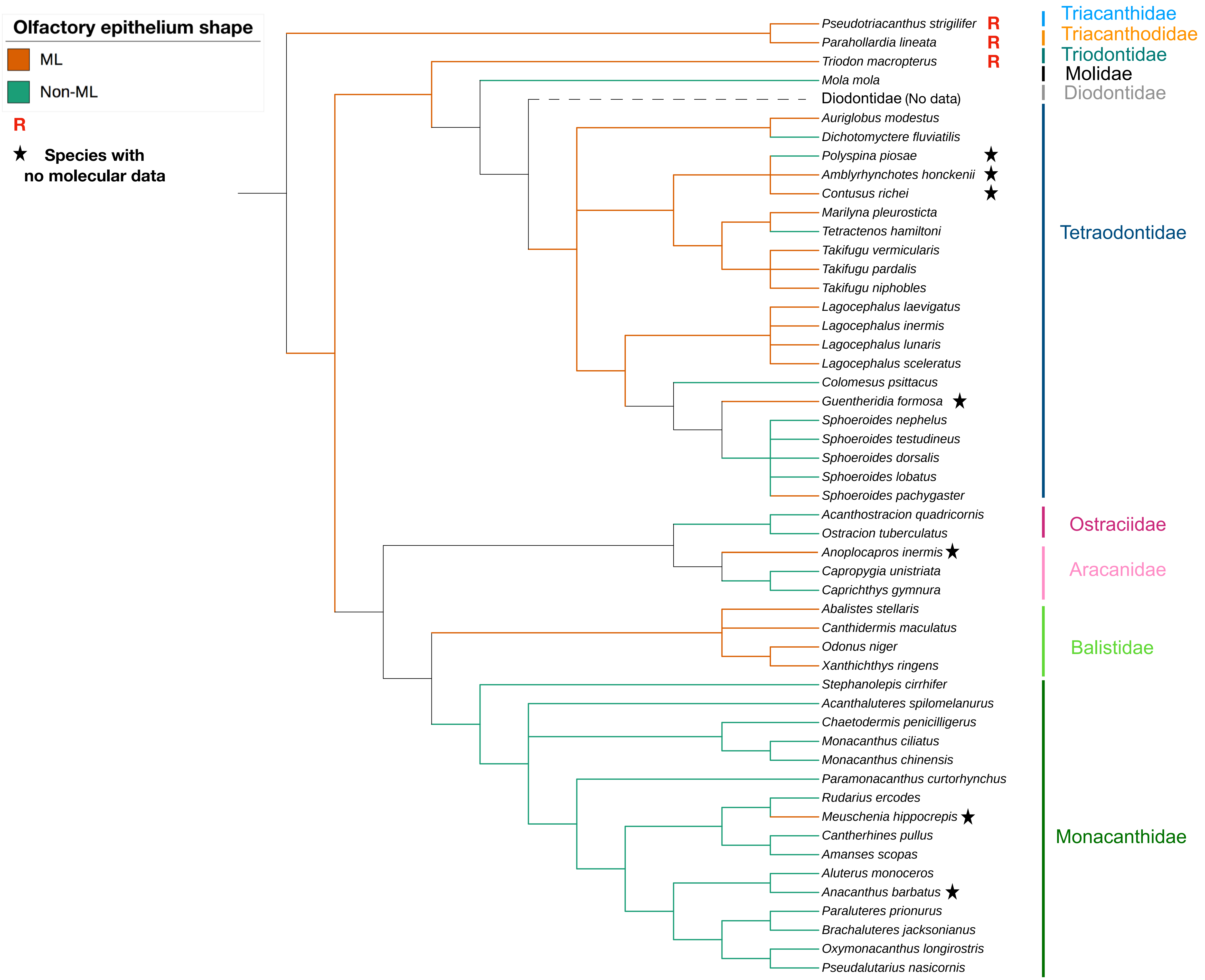

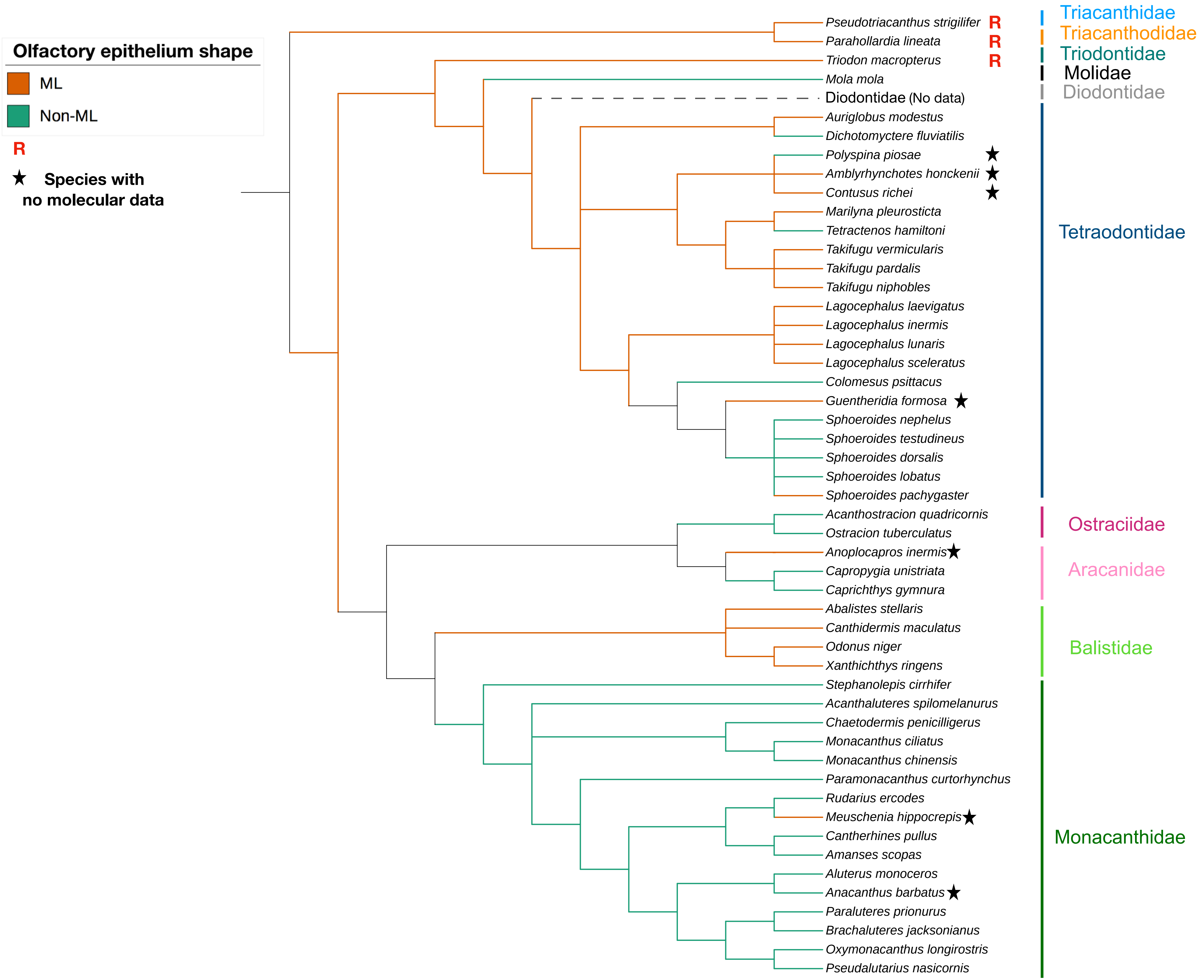

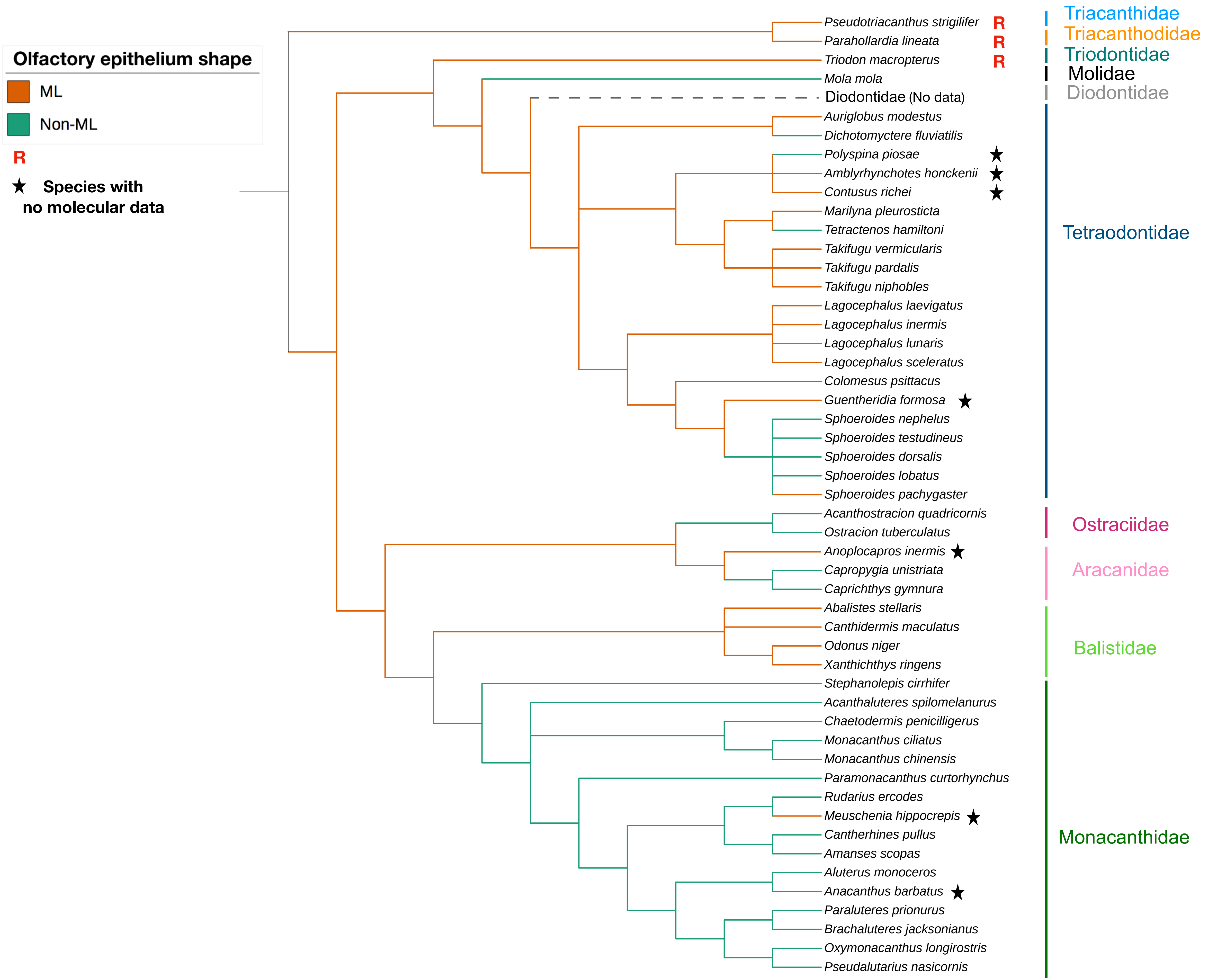

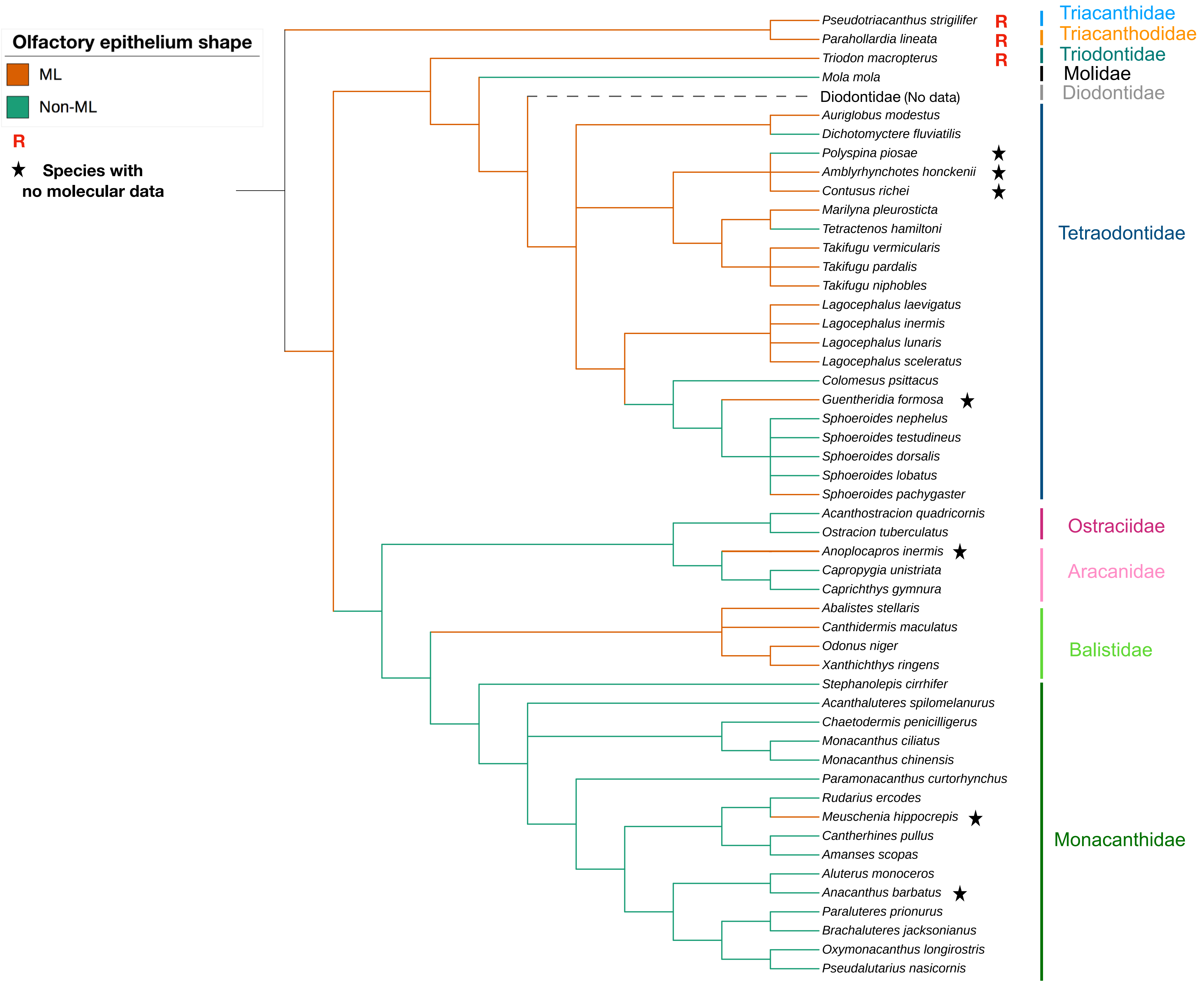

### Supplementary fig. S12

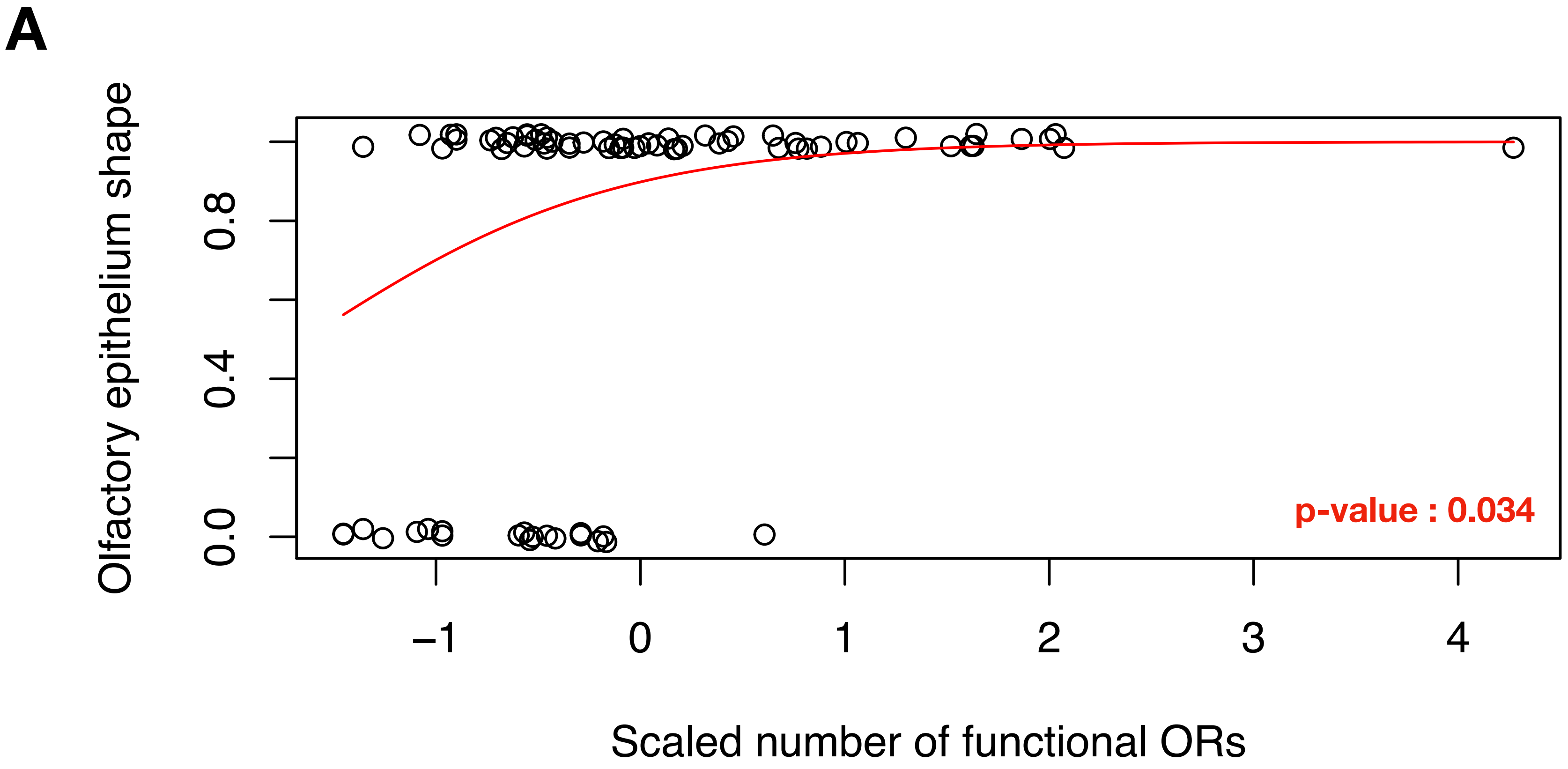

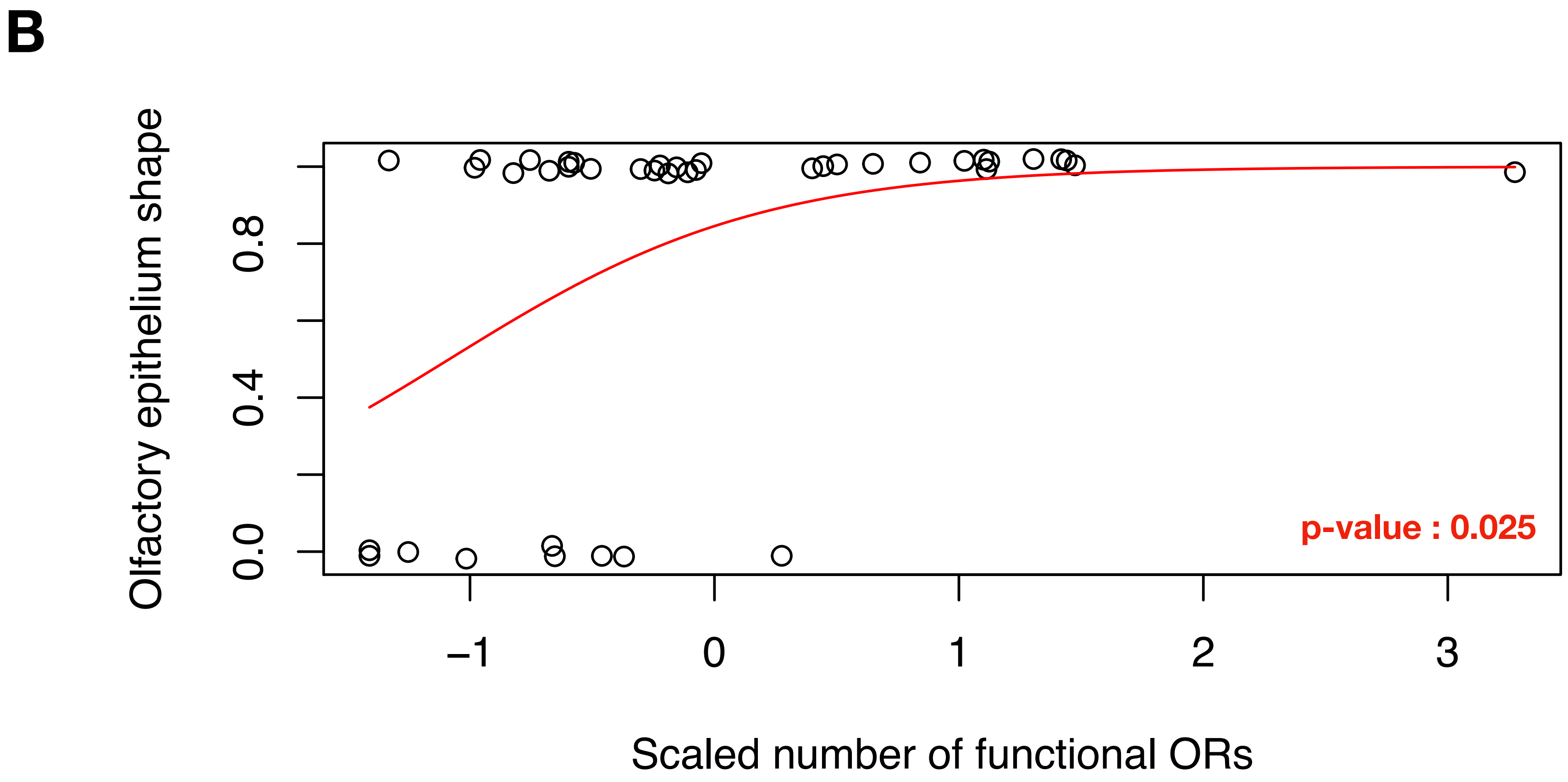

### Supplementary fig. S13

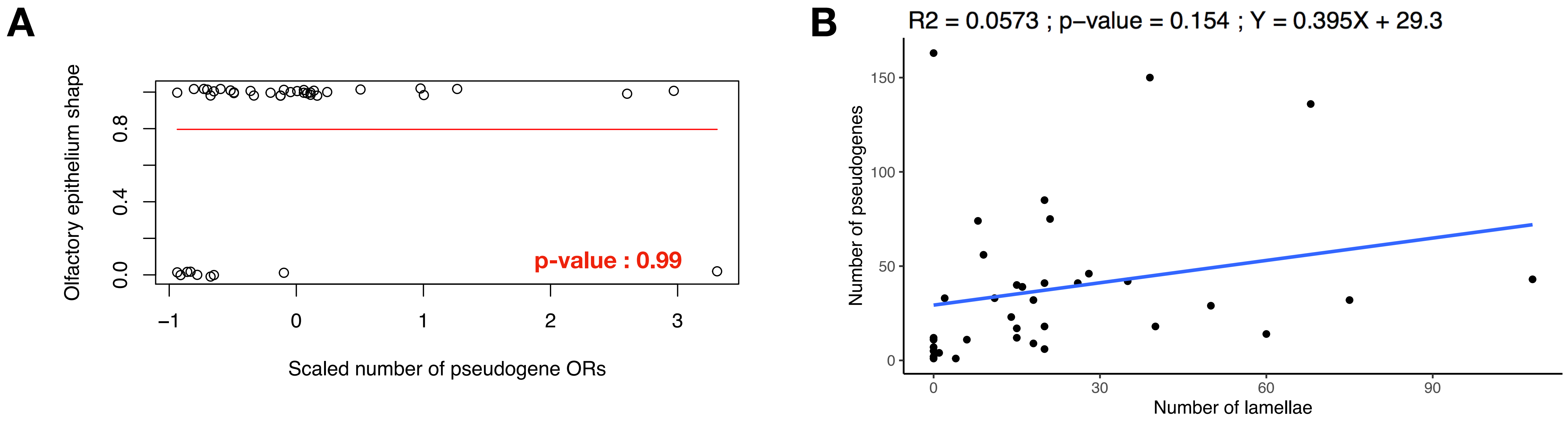

### Supplementary fig. S14

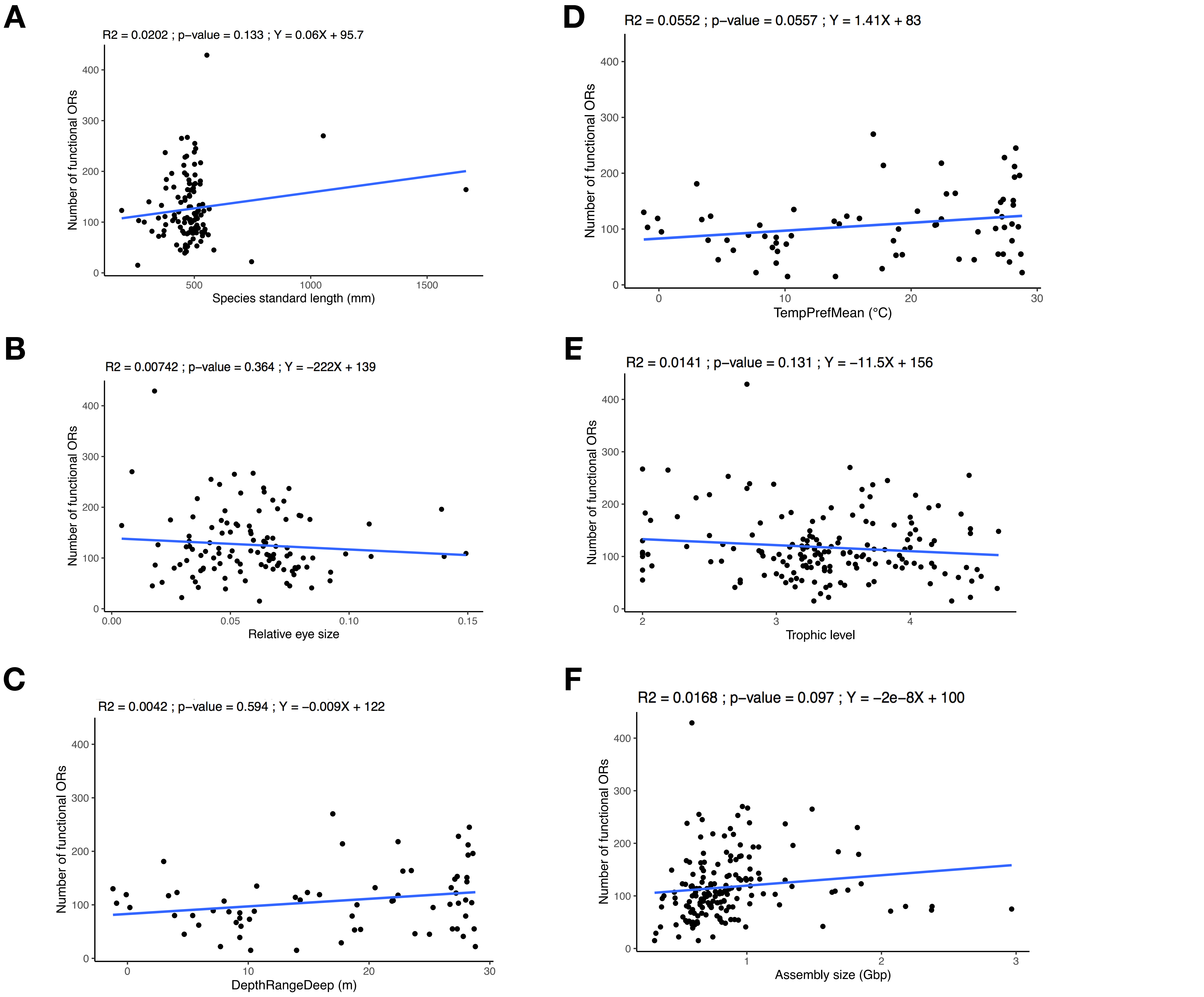
